# Validation of optogenetic approach to investigate fatigable weakness using a zebrafish model of *syt2* congenital myasthenic syndrome

**DOI:** 10.64898/2026.05.30.728923

**Authors:** J. M. G. Lau, S. F. Gaudreau, K. K. Palacek, S. Spendiff, T. V. Bui, H. Lochmüller

## Abstract

Congenital myasthenic syndromes (CMS) are rare inherited diseases of the neuromuscular junction (NMJ). There are 40 identified CMS genes, but many patients go without genetic diagnosis, which suggests new genes have yet to be discovered and characterised. Here, we describe an optogenetic approach to study fatigable muscle weakness and NMJ function in larval zebrafish to facilitate screening approaches for uncovering novel CMS genes. Using blue-light illumination of spinal motoneurons that express channelrhodopsin-2 (ChR2) to induce muscle contraction, we measure motor defects at the behavioural, synaptic, and genetic level through a novel behavioural assay, standard whole-cell electrophysiology of individual muscle fibers and a customized NMJ gene panel. We employ this approach in synaptotagmin-2 (*syt2*) morphant zebrafish, an identified CMS gene model, to validate its usefulness. Our customized optogenetic behavioural assay successfully demonstrates reduced, fatigable, locomotor response during repeated activation of spinal motoneurons. Whole-cell electrophysiology recordings of optogenetically-elicited endplate currents in muscle fibers reveal similarities to altered properties of NMJ function in *syt2* morphants reported in other studies using the standard paired motoneuron-muscle electrophysiology technique. Finally, we develop a genetic panel of CMS and NMJ-related genes to characterize the expression landscape of *syt2* morphants to elucidate potential pathomechanisms and novel therapeutic targets. We propose that this three-tiered approach successfully links behaviour, synaptic motor function, and genetic expression and can be used as a tool in the screening of novel genes associated with CMS.

## Introduction

Congenital myasthenic syndromes (CMS) are a diverse group of rare inherited neuromuscular disorders. CMS onset typically presents at birth or early childhood but can also appear in adolescence (1–3). The defining symptoms include fatigable muscle weakness, ptosis, and decremental response during repetitive nerve stimulation electromyography (RNS-EMG) (1). The root cause of CMS are mutations in genes with relation to the formation, function, and maintenance of the neuromuscular junction (NMJ). Currently, there are about 40 genes that have been linked to CMS and are extensively described (1,2). The most common forms of CMS affect subunits of the AChR and lead to reduced numbers of AChRs at the motor endplate, modify ACh binding, or alter AChR channel kinetics (1–5). However, even with 40 known CMS genes, about 20% of CMS patients go without definitive genetic diagnosis, indicating that novel genes have yet to be identified (6).

Larval zebrafish are an attractive platform for screening novel neuromuscular gene candidates. First, their high fecundity and rapid development render them favourable to screening studies. Next, their genetic amenability permits efficient generation of CRISPR-Cas9 or morpholino (MO) mediated gene knock-outs and knock-downs respectively (7–11) and the rapid development of their motor systems, as early as 17 hours post-fertilization (hpf), allows for quick phenotype characterization (12). Zebrafish also have well-defined motor movements, which emerge within the first day of development, and ultimately reach mature larval swimming by 4 days post-fertilization (dpf) which allows for standardized measurements (12–15). Additionally, orthologous genes associated with the NMJ share common functions across vertebrates, highlighting that key functional architecture of the zebrafish NMJ is similar to humans (13–15). These advantageous features of larval zebrafish facilitate high throughput loss-of-function testing of NMJ proteins. Indeed, larval zebrafish have been used to generate numerous models of neuromuscular diseases (NMD) including muscular dystrophies (13,16), amyotrophic lateral sclerosis (ALS) (17,18), spinal muscular atrophy (SMA) ((14)), and CMS among others (14,19–23).

Accurate validation of novel CMS gene candidates would ideally include assessment of locomotor behaviour as well as NMJ function when using *in vivo* models such as zebrafish (24,25). With zebrafish, typical motor assays include observing chorion movement of embryos, the touch-evoked escape response, as well as activity of free-swimming larvae (12,26). A well-established method for assessing neural activity across the NMJ in larval zebrafish involves a paired patch-clamping method targeting a primary motoneuron and corresponding fast muscle cell simultaneously (27). Endplate currents (EPCs) at the level of the skeletal muscle are recorded in voltage-clamp configuration during electrical stimulation of the motoneuron (27). Using this technique, effects on NMJ signal transmission have revealed the roles of numerous NMJ proteins, shedding light onto fundamental physiological properties of signal transmission at the NMJ of larval zebrafish (19,28–31).

Here, we propose a technically less challenging approach to screen for novel CMS gene candidates with robust behavioral, electrophysiological, and NMJ-gene expression data. Our approach utilizes a transgenic zebrafish line, 1020:UAS;Gal4:ChR2(H134)-mCherry (ChR2-mCherry) (32,33)wherein blue light activated channelrhodopsin-2 (ChR2) is expressed in spinal interneurons as well as both primary and secondary motoneurons. Excitation of these neurons by blue light elicits swimming in larval zebrafish (32). Indeed, this line has been used previously to excite motoneurons while performing patch-clamp electrophysiology in skeletal muscle to investigate calcium channel contributions to NMJ signal transmission in zebrafish(28). By circumventing the need to attach individual electrodes to both a motoneuron and its paired, innervated muscle fiber, optogenetic stimulation in this transgenic zebrafish line can be used to easily excite motoneurons for investigations into both locomotor and NMJ function. Using the previously identified CMS gene synaptotagmin-2 (*syt2*) (34) to generate knockdowns in ChR2-mCherry embryos, we demonstrate that this optogenetic approach permits facile assessment of movement and NMJ function using blue-light illumination. Additionally, our NMJ-gene panel adds an additional dimension to interpreting behavioral and electrophysiological results and reveals further lines of investigation for CMS pathomechanisms. Not only is this screening procedure technically more approachable but also provides a behaviourally relevant assessment of NMJ function. We highlight the benefits of using this approach as a tool for screening CMS genes, all the while underscoring its limitations.

## Methods

### Zebrafish husbandry

Adult and larval zebrafish of AB lineage were maintained in accordance with the University of Ottawa’s Animal Care and Veterinary Services (Protocol numbers: CHEOb-4195, CHEOe-4191, BLe-4416). In brief, adult fish were kept at 28°C on a 14 h/10 h light/dark cycle in continuous flow tanks at pH 6.8 - 7.6. The night before breeding, 1 male and 1 female were placed in a breeding trap separated with a transparent divider. The following morning, the dividers were removed and the fish allowed to breed. Eggs were collected from the bottom of the trap and immediately placed in warmed E3 medium.

### Morpholino microinjections

Antisense morpholino oligonucleotides (MOs) were designed and synthesized by Gene Tools, LLC (Philomath, OR, USA) targeting the *syt2* translation start site (*syt2* MO) with the sequence: 5’-AGCCAGAGATCATGAAGTGGAACCT-3’ as found in previous literature (19). A standard control morpholino against human beta-globulin, with no known zebrafish targets, was used as a negative control. Stock aliquots of MOs were resuspended in nuclease-free water to 3 mM and stored at −20°C. The night before injection, MO working solutions were prepared with nuclease-free water to a volume of 9 µL. Immediately prior to injection, 1 µL 0.05% phenol red was added as a tracer for a final MO concentration of 1 mM. Embryos were then collected within 5 min post-fertilization and aligned on a 2% agarose injection mold. MO solutions were loaded into pulled borosilicate glass capillary needles and injected into the yolk of 1-4 cell stage embryos using a Narishige IM-300 microinjector. Injection volume was calibrated to deliver up to 6 ng per injection, ensuring that droplet volume did not exceed 10% of yolk volume. Injected embryos were transferred to fresh E3 medium and incubated at 28.5°C in a dark incubator for further development.

### Whole-mount immunofluorescent staining

Immunofluorescent staining was performed directly on 1020:Gal4;UAS:ChR2(H134)-mCherry (herein referred to as ChR2-mCherry) zebrafish without clearing agents since this line lacks pigmentation. In brief, euthanized 3 dpf embryos were fixed in 4% paraformaldehyde (PFA) in phosphate buffered saline (PBS) overnight. Embryos were then washed 3x with PBS, then treated with collagenase A (Millipore Sigma, 1 mg/ml) for 30 min at room temperature (RT) and washed 3x for 5 min with PBS. Embryos were then permeabilized for 7 min in 100% ice-cold acetone, then washed 3x for 5 min in PBS. Embryos were then blocked with 5% horse serum (HS) in PBS with Tween 20 (PBST) at RT for 1 hour. After blocking, embryos were incubated overnight at 4°C with primary antibody on an orbital shaker (DSHB, SV2, 1:500; DSHB, ZNP-1, 1:500). The following day, embryos were washed 5x for 20 min in PBST, then incubated with α-Bungarotoxin Alexa FluorTM conjugate (Invitrogen B13422, 1:1000) and 594 Alexa FluorTM goat anti-mouse IgG1 antibody (Invitrogen A21125, 1:200) for 2 hours at RT. Embryos were then washed 2x for 5 min, then overnight in PBST at 4°C. Fish were the mounted and imaged using 5 µm Z-stacks on an Olympus Fluoview FV-1000 Laser Confocal Microscope under the 20X or 40X objective lens.

### Preparation for electrophysiology

Larval ChR2-mCherry (32,33) zebrafish aged 2-3 dpf were prepared for electrophysiology similarly to as is described previously (27). Larvae were anesthetized in 0.02% tricaine (MS-222, Aqualife TMS; Syndel Laboratories) before being pinned down through the notochord onto a Sylgard (Dow Corning) coated dish. Pins were made using tungsten wire (diameter of 0.025 mm); one was placed caudally near the tip of the tail, and the other was placed rostrally near the center of the yolk sac. Spinalization was performed using fine surgical scissors at the level of somites 2-3. The skin was then peeled back between the two pins using fine forceps. Next, larvae were bathed in a 2M formamide solution (Millipore Sigma, F9037; 2-5 min). The formamide solution was rinsed out 4-5 times with extracellular recording solution (see below). Superficial slow muscle fiber removal was performed using suction through a wide-bored glass capillary.

Muscles overlaying the spinal cord were not removed.

### Electrophysiology

Extracellular recording solution consisted of artificial cerebrospinal fluid (aCSF) containing: 134 mM NaCl, 2.9 mM KCl, 1.2 mM MgCl_2_, 2.1 mM CaCl_2_, 10 mM dextrose, and 10 mM HEPES, (pH of 7.8 adjusted with NaOH and osmolarity between 280-290 adjusted with sucrose).

Borosilicate glass capillaries (outer diameter: 1.5 mm; inner diameter: 1.1 mm, Sutter Instruments, catalog #BF150-110-10) were used to form pipette tips having a resistance between 2-3 MΩ. These microelectrodes were backfilled with intracellular recording solution consisting of: 116 mM potassium gluconate, 16 mM KCl, 2 mM MgCl_2_, 10 mM HEPES, 10 mM EGTA, and 4 mM Na_2_ATP, osmolarity adjusted to 290 mOsm, pH 7.2 adjusted with KOH (35). Ventral fast muscle fibers innervated by caudal primary (CaP) motoneurons were targeted for patch-clamp electrophysiology. Once a giga-ohm seal was formed, light suction was applied to break into the membrane of the muscle fiber. As described previously (27), membrane potential was held at -50 mV for the duration of the recording. To stimulate motoneurons optogenetically, blue light (480 nm) illumination was applied over an area of the spinal cord corresponding to approximately one spinal segment that contains the CaP motoneuron innervating ventral fast muscle fibers. Individual light pulses were delivered at a duration of 5 ms. The signal was recorded in voltage-clamp mode, amplified and sampled at 10 kHz with a Multiclamp 700B from Axon Instruments (Molecular Devices) and finally digitized with a Digidata 1550 (Molecular Devices). Recordings were made from mid-body somites located within 4 somites rostral and 4 somites caudal to the anal pore.

### Gene panel design

We utilized Search Tool for Recurring Instances of Neighbouring Genes (STRING version 11) and Ingenuity Pathway Analysis (IPA, Qiagen) to generate interaction networks based on the known CMS genes *AGRN, ALG14, ALG2, CHAT, CHD8, CHRNA1, CHRNB1, CHRND, CHRNE, CHRNG, COL13A1, COLQ, DOK7, DPAGT1, GFPT1, GMPPB, LAMA5, LAMB2, LRP4, MUSK, MYO9A, PLEC, PREPL, PURA, RAPSN, RPH3A, SCN4A, SLC18A3, SLC25A1, SLC5A7, SNAP25, SYT2, TOR1AIP1, UNC13A, VAMP1,* and *MACF1*. This mapping was repeated based on human, mouse, and zebrafish datasets and extended to include the top 200 nodes. We then cross-referenced these generated gene interaction networks with RNA-seq lists from Ham *et al*., 2025 and Hui et al, 2021 to identify genes present in at least two of the lists (36,37). This list was submitted to NanoString (Bruker Spatial Biology, USA) for panel design using their barcoding fluorophore technology. Pooled zebrafish RNA was loaded on to an nCounter® Analysis System (NanoString Technologies, Seattle, WA, USA) for transcript analysis. A full list of gene targets, probe sequences, and reference genes can be found in **Supplemental 1 Appendix**.

### RNA Extraction and running of NanoString CMS panel

RNA extraction was performed on pooled samples of a maximum of 30 zebrafish per sample. Extractions were performed using the Qiagen RNeasy Fibrous Tissue Kit (74704) and samples were stored at -80°C or used directly for hybridization and running of the panel according to the manufacturer’s protocol.

### Locomotor studies

#### Chorion activity recording and analysis –

At 24 hpf, larvae were placed in a 10 cm petri dish with just enough E3 medium to keep them wet and ensure they sat stationary on the bottom of the dish to prevent drifting during recording. They were then placed on the stage of a Leica EZ4 W stereomicroscope and recorded at 30 frames per second (fps) for 1 min. Activity was then analyzed using Danioscope software (Noldus, Netherlands).

#### Touch-evoked response –

At 2 dpf embryos were removed from their chorion using fine forceps and individually placed into a 10 cm petri dish on nitrocellulose paper to ensure good contrast. Each fish was then stimulated on the back of the head with a fine pipette tip, and the response was recorded using a Sony highspeed camera (Sony RXO II) at 240 fps.

#### Light/dark transition assay –

Light/dark transition assays were preformed between 5 to 7 dpf where fish were placed in a 48-well plate with 300 µL E3 medium immediately before recording. Plates of fish were placed in a Zantiks MWP automated tracking system (Cambridge, UK) and allowed to acclimate for 5 min. Sequentially, fish were then subject to 3 cycles of 300 s white light, then 300 s of darkness. The internal temperature of the system was maintained at 28°C throughout the recording.

#### Fatigable weakness assay –

This assay was performed at 7 dpf when ChR2-mCherry zebrafish larvae showed sufficient behavioral response to blue LED light (450 nm) stimulation. This protocol consisted of four blocks of twenty 1-s pulses of full intensity blue light stimulation applied at a constant inter-pulse interval with decreasing inter-pulse interval with each successive recovery block (60 s, 30 s, 15 s, 5 s). Each recovery block was separated by a 5 min rest period.

### Statistical analysis

Statistical analysis and plotting of immunofluorescence images, morphological features, chorion activity, touch response assay, light-dark assay, and fatigable weakness assay is specified in the figure captions and were chosen based on data distribution and experimental design. For analysis of fatigable weakness data, only distance data during and for the first 5 s post blue light stimulus were analyzed to account for the shortest rest interval. All electrophysiological data was saved as .abf files. We used the open source pyABF python package to import and read .abf files in Spyder (version 5.1.5). Analysis of recordings was semi-automated using Python (version 3.9.12) to detect spontaneous miniature end-plate currents (mEPCs) as well as evoked end-plate currents (EPCs). Latency of EPCs was measured as the time between the onset of the blue-light pulse and the peak of the EPC. Quantal content, number of release sites, release probability, and levels of steady-state depression were estimated as described previously (21). Assessment of the contributions of synchronous and asynchronous release at the NMJ was performed similarly to Wen *et al*. (19). The time integral of synaptic current was computed to quantify charge transfer at the NMJ. Synaptic events were classified as synchronous if occurring within the first 15 ms from the onset of the blue-light pulse and asynchronous if occurring during the remaining 35 ms of the 50-ms interpulse interval of the 20 Hz train. Data are reported as mean ± SD. Given the known mosaicism and variability of morpholino-mediated knockdown, outlier analysis was performed using the ROUT method (Q = 1%). This was applied uniformly across all datasets. All statistical tests employed are specified in the figure captions. All violin plots represent the median with a solid black line and the 1st and 3rd quartiles as black dashed lines. Gene panel analysis was completed using pipelines in nSolver® Analysis Software version 4.0 (Bruker Spatial Biology, USA).

## Results

### Translation-blocking syt2 MO inhibits expression of syt2 and increases myotomal AChR clustering

To first validate the *syt2* translation-blocking MO from Wen *et al*. (19), we injected WT zebrafish eggs at the 1-2 cell stage with 6 ng of MO. To confirm knockdown of *syt2* at the NMJ in zebrafish we performed immunofluorescent staining at 3 dpf with the *syt2-*specific antibody ZNP-1 and 488 alexa fluor conjugated α-bungarotoxin to label post-synaptic AChRs (**Fig. 1*A***). Firstly, upon *syt2* knockdown, general morphological characteristics showed slight decreases in body length and eye size with no significant signs of off-target effects such as cardiac pulmonary edema (data not shown) (38). AChR colocalization with syt2, as measured by Pearson’s coefficient, was significantly decreased when compared to controls, indicating successful knockdown of syt2 (**Fig. 1*B***; uninjected = 0.586 ± 0.0437; control MO = 0.594 ± 0.0292; *syt2* MO = 0.153 ± 0.0462). Additionally, overall myotome area was significantly decreased and indicates amount of muscle tissue is reduced in *syt2* morphants (**Fig. 1*C***; uninjected = 6,670 ± 671 µm2; control MO = 6,150 ± 682 µm2; *syt2* MO = 4010 ± 683 µm2). To account for this difference in area, we measured the density and average area of both AChR and ZNP-1-positive clusters in each myotome. We observed an increase in myotomal AChR spot density in *syt2* morphants (**Fig. 1*D***; uninjected = 8.65 x 10-3 ± 2.25 × 10-3 spots/µm2; control MO = 6.87 × 10-3 ± 2.63 × 10-3 spots/µm2; *syt2* MO = 1.17 × 10-2 ± 3.71 × 10-3 spots/µm2) but no change in average area of these AChR clusters (**Fig. 1*E***; uninjected = 4.79 ± 1.39 µm2; control MO = 4.58 ± 0.719 µm2; *syt2* MO = 4.77 ± 0.232 µm2). Increased number of AChR spots could indicate that AChR pre-patterning is also increased. Analysis of ZNP-1 positive spots shows typical decreases in both density (**Fig. 1*F***; uninjected = 1.12 × 10-2 ± 3.73 × 10-3 spots/µm2; control MO = 8.34 × 10-3 ± 4.39 × 10-3 spots/µm2; *syt2* MO = 5.23 × 10-4 ± 3.49 × 10-4 spots/µm2) and average area (**Fig. 1*G***; uninjected = 4.69 ± 1.00 µm2; control MO = 3.19 ± 0.694 µm2; *syt2* MO = 1.40 ± 0.468 µm2), indicative of successful *syt2* knockdown. Both Mander’s M1 and M2 were calculated to observe the proportions of myotomal AChR spots colocalized with syt2 spots and vice versa, both of which were decreased in *syt2* morphants due to the lack of syt2 protein (**Fig. S2*B-C***).

**Figure 1.**
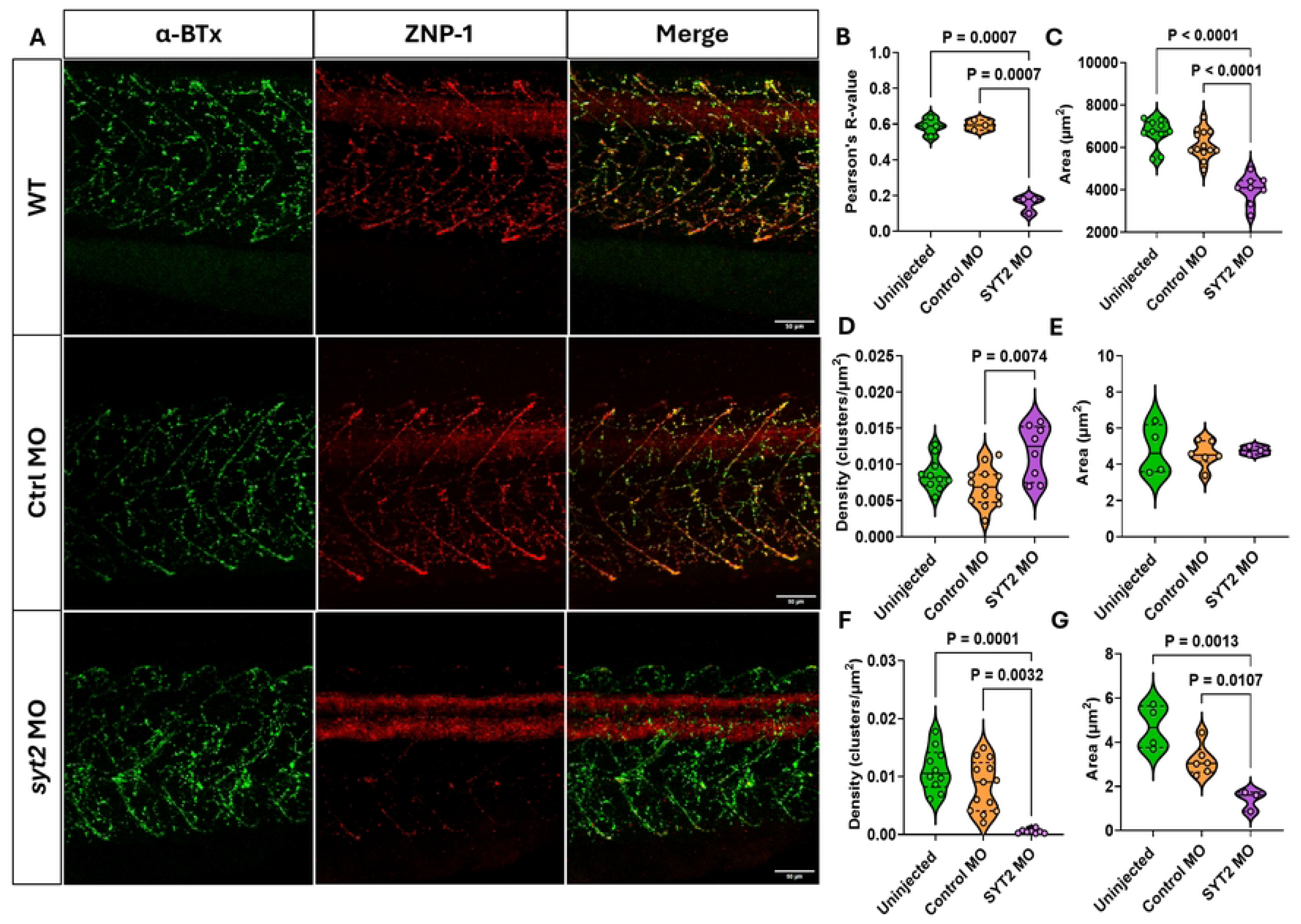
Immunofluorescent analysis of *syt2* expression at the neuromuscular junction in *syt2* morphants. **(A)** Representative immunofluorescent staining of NMJ in 3 days post fertilization (dpf) of uninjected (WT, *n =* 3), control morpholino injected (control MO,6 ng, *n* = 4), and *syt2* MO injected (*syt2* MO, 6 ng, *n* = 3) zebrafish larvae (N = 2-6 myotomes/larvae). Acetylcholine receptors (AChRs) are stained with alpha-bungarotoxin (green, first column) and *syt2* with ZNP-1 (red, middle column), merged images of red and green channels show location of neuromuscular junctions (yellow). Scale bar = 50 µm. **(B)** Quantification of colocalized myotomal signal, excluding the spinal cord, showing that *syt2* morphants have reduced Syt2 colocalization with AChRs. **(C)** Measured myotome area of *syt2* morphants compared to controls shows decreased myotomal size in *syt2* morphants. **(D)** Density of myotomal postsynaptic AChR spots are increased in *syt2* morphants but **(E)** AChR cluster size does not change as compared to uninjected and control morphant larvae. **(F)** The myotomal density of ZNP-1 positive spots and their **(G)** average size are both significantly reduced in *syt2* morphants when compared to controls, suggesting successful knockdown of *syt2*. Scale bar = 50 µm. *Statistical analysis*, 2-way ANOVA with Tukey’s multiple comparison’s test.

### Frequency of miniature end-plate currents increased in syt2 morphant fish

We chose to generate *syt2* morpholino (MO) knock-down using the ChR2-mCherry line specifically since NMJ function in *syt2* morphants has already been described (19). This has facilitated direct comparisons with Wen et al. (2010)’s data to validate our proposed optogenetic approach to assess NMJ function. A key characteristic of NMJ function in *syt2* MO fish is the increased frequency of spontaneous miniature mEPCs recorded in fast muscle fibers innervated by CaP motoneurons, as described previously (27). In assessing the frequency of mEPCs (**Fig. 2*A-*C**), we found that the number of spontaneous mEPCs generated per s in ventral fast muscle fibers in our *syt2* morphants increased 16 (vs. control MO) to 29 (vs. uninjected)-fold compared that of controls (**Fig. 2*D***; uninjected control: 0.0539 ± 0.0387 events/s; control MO: 0.0985 ± 0.0730 events/s; *syt2* MO : 1.62 ± 1.28 events/s). These data help confirm successful knock-down of *syt2* in our *syt2* morphants.

**Figure 2.**
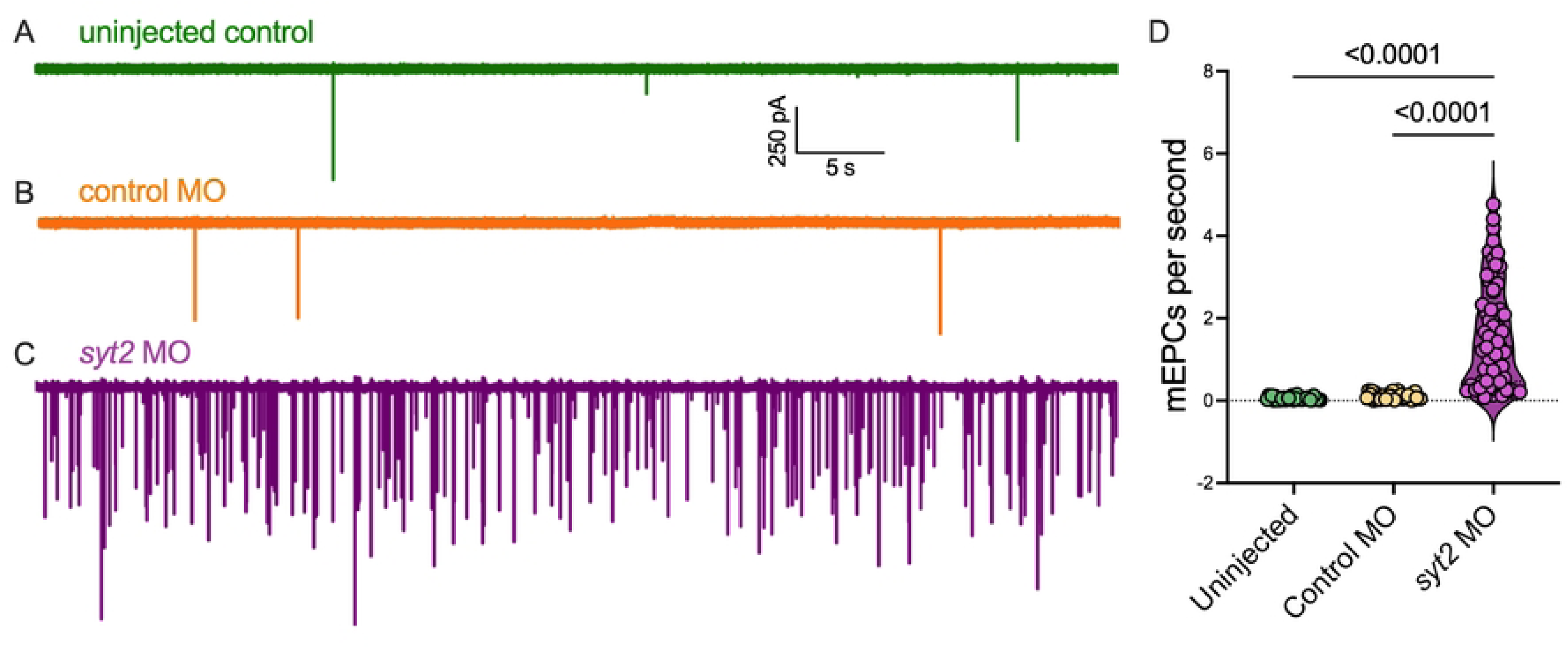
Increased spontaneous miniature end-plate currents in *syt2* MO fish. **(A)** Representative trace of spontaneous mEPCs recorded from a ventral fast muscle fiber of an uninjected control fish (*n* = 33 muscle fibers; *N* = 19 animals). **(B)** Representative trace of spontaneous mEPCs recorded from a ventral fast muscle fiber of a control MO fish (*n* = 32; *N* = 16). **(C)** Representative trace of spontaneous mEPCs recorded from a ventral fast muscle fiber of a *syt2* MO fish (*n* = 59; *N*= 28). **(D)** Number of spontaneous mEPCs recorded per second (uninjected versus control MO: P = 0.3689; uninjected versus *syt2* MO: P < 0.0001; control MO versus *syt2* MO: P < 0.0001). *Statistical analysis*, Kruskal-Wallis test with Dunn’s multiple comparisons test.

### syt2 morphants exhibit locomotor development defects in chorion activity, touch-response, and light-dark transition test

We then characterized locomotor ability of *syt2* morphants. The first measurable behavior of locomotor development in zebrafish are the spontaneous coiling/tumbling movements of zebrafish larvae in their eggs (12). To measure this activity, we recorded the embryos in their chorions at 24 hpf for 1 min and found that, as compared to uninjected and control MO injected clutch mates, the frequency of movement in *syt2* morphants was significantly impaired (**Fig. 3*A***; uninjected = 0.293 ± 0.184 Hz; control MO = 0.337 ± 0.208 Hz; *syt2* MO = 0.244 ± 0.224 Hz). Additionally, we performed a touch-response assay to measure the average velocity of larvae after physical stimulus by touching a pipette tip to the base of the skull at 2 dpf. The average velocity of this response was also seen to be decreased in *syt2* morphants (**Fig. 3*B***; uninjected = 100 ± 26.3 %; control MO = 93.3 ± 18.5 %; *syt2* MO = 69.4 ± 17.1 %). Reductions in the velocity of this response signal muscle fatigability in *syt2* morphant larvae as there is no significant decrease in the overall distance travelled after stimulus (**Fig. S2*D***; uninjected = 100 ± 48.9 %; control MO = 82.4 ± 32.1 %; *syt2* MO = 57.1 ± 30.7 %) or the strength of acceleration upon stimulus (**Fig. S2*E***; uninjected = 100 ± 37.8 %; control MO = 115 ± 45.5%; *syt2* MO = 117 ± 43.6 %). Lastly, we measured the movement of *syt2* morphants at 5 dpf using the light-dark transition assay where larvae are exposed to 5 min of alternating light/dark cycles after a 10 min acclimation (**Fig. 3*E***). When analyzing total activity (**Fig. 3*C***; uninjected = 100 ± 77.2 %; control MO = 97.5 ± 77.9 %; *syt2* MO = 9.52 ± 9.23 %) and dark phase activity (**Fig. 3*D***; uninjected = 100 ± 82.5 %; control MO = 112 ± 90.4 %; *syt2* MO = 8.55 ± 8.61 %), we can see that while *syt2* morphants still respond to dark stimulus, where most movement occurs, they move ∼10-fold less than their uninjected and control MO injected clutch mates.

**Figure 3.**
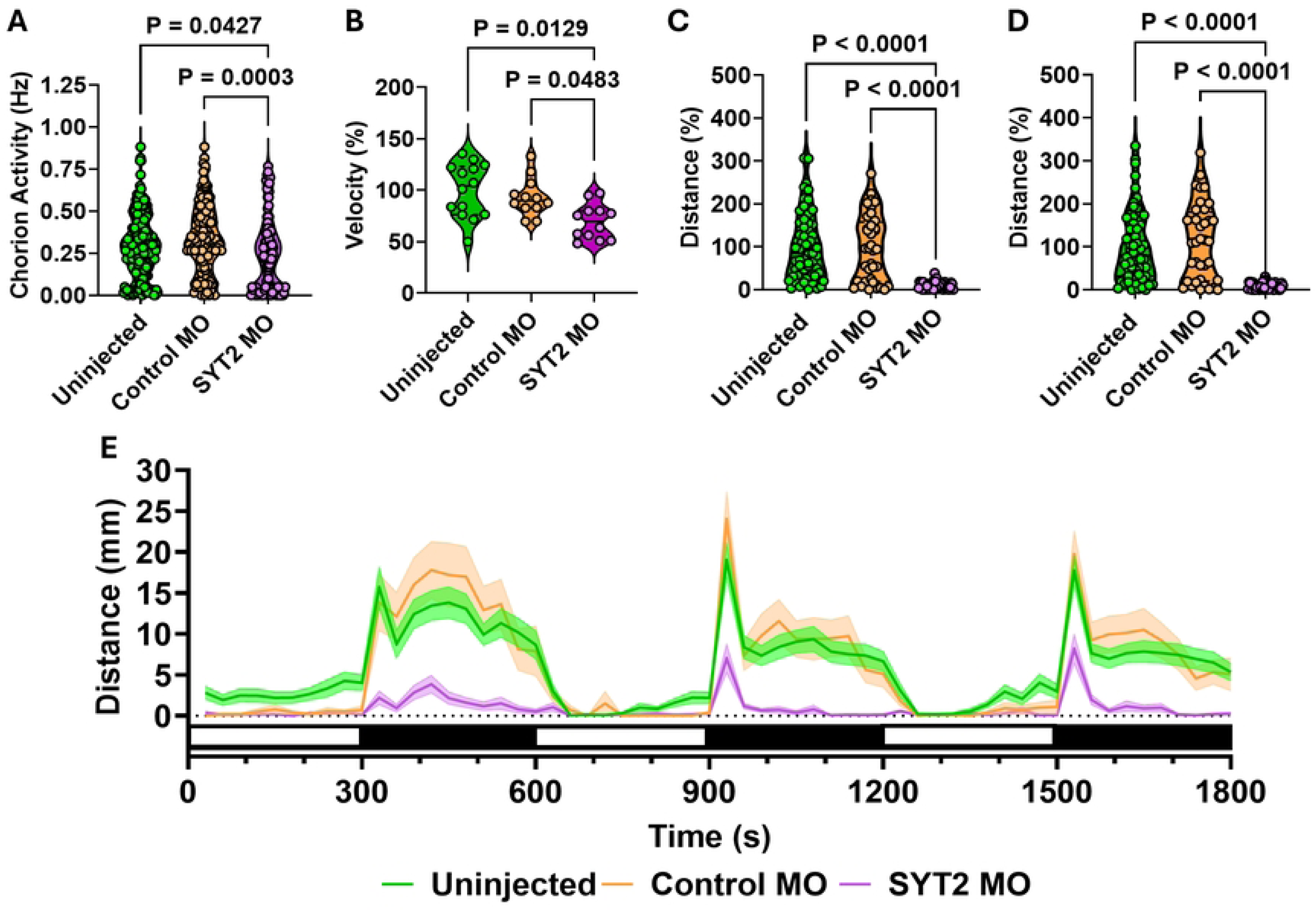
Early assessment of milestone developmental locomotor behaviors in s*yt2* morphants. **(A)** Chorion activity at 24 hpf, showing activity in *syt2* morphants is decreased compared to controls (uninjected, *n* = 204 fish; Ctrl MO, *n* = 120; SYT2 MO, *n* = 103). **(B)** A significant decrease in the average velocity during the touch-evoked response assay in 3 dpf *syt2* morphants was observed when compared to controls (uninjected, *n* = 14; Ctrl MO, *n* = 12; SYT2 MO, *n* = 12). **(C)** Total distance travelled during the Light-dark transition test (LDT) where 5 dpf larvae were acclimated for 10 min (not recorded), then subjected to 3 cycles of alternating 5 min light/dark stimulus at 28° C (uninjected, *n* = 61; Ctrl MO, *n* = 36; SYT2 MO, *n* = 48). **(D)** Distance travelled during the dark phase of the LDT test. **(E)** Overall trace of the light (white bars)-dark (black bars) transition test. *Statistical analysis*, Kruskal-Wallis test with Dunn’s multiple comparisons test.

### syt2 is required for response at high frequency stimulation in optogenetic zebrafish line

To test this optogenetic model’s ability to respond to blue light stimulation and to define skeletal muscle fatigability, we designed our fatigability assay to mimic a basic, functional test which CMS patients may be asked to perform. During this test patients raise their arms 10 times to the front or side in succession, allowed to recover, then asked to repeat the procedure (39). In classical cases CMS patients fail to raise their arms after the 2nd or 3rd try as NMJ transmission defects begin to present and muscle contraction fails. However, after recovery, patients regain the ability to raise their arms, showing that NMJ signalling can be restored upon rest. To mimic this, we took advantage of the optogenetic zebrafish line ChR2-mCherry to precisely evoke a swim response in zebrafish larvae at 7 dpf (32). This optogenetic fatigability assay consisted of 4 recovery blocks with interspersed blue light stimulation. The first block consisted of twenty 1 s long, blue light pulses with a constant inter-pulse interval, which were followed by 5 min rest periods. The inter-pulse intervals of each successive recovery block decreased from 60 s to 30 s, 15 s and 5 s to provide less recovery time between stimulation pulses. Each block was separated by a dark rest period of 5 min without stimulation (**Fig. 4*A***). When *syt2* morphants were subjected to this optogenetic fatigability assay, we observed a significant drop in their overall response (evoked response) as compared to uninjected and control MO injected clutch mates in all recovery blocks (**Fig. 4*B***). Interestingly, when comparing evoked swim response within control groups (uninjected and control MO injected) we can see a decline in response from block to block. Eventually the decline in block-to-block response becomes significant when comparing all other recovery blocks to the 5 s recovery block, suggesting this point is where 7 dpf fish larvae become naturally fatigued (**Fig. S2*G***). Total decline in response from 60 s to 5 s was 18.9 ± 3.06 % in uninjected larvae, 15.5 ± 3.02 % in control MO morphants, and 11.0 ± 2.30 % *syt2* morphants (**Fig. S2*G***). Although *syt2* morphants had the least decline in evoked swim response, we note that *syt2* morphants showed the lowest response across all recovery blocks and approaches zero overall. In responsive fish, which we defined as fish with positive acceleration during optogenetic stimulus, we also measured distance travelled after stimulus, average velocity, and average evoked acceleration across all recovery blocks to identify trends in these metrics. Distance was defined as total distance travelled during and for 5 s post stimulus to account for the smallest duration of recovery given to the fish. Velocity was defined as the average speed over the same time span as defined above. Evoked acceleration was measured as an increase in velocity from the last second pre-stimulus to the first second during stimulus.

**Figure 4.**
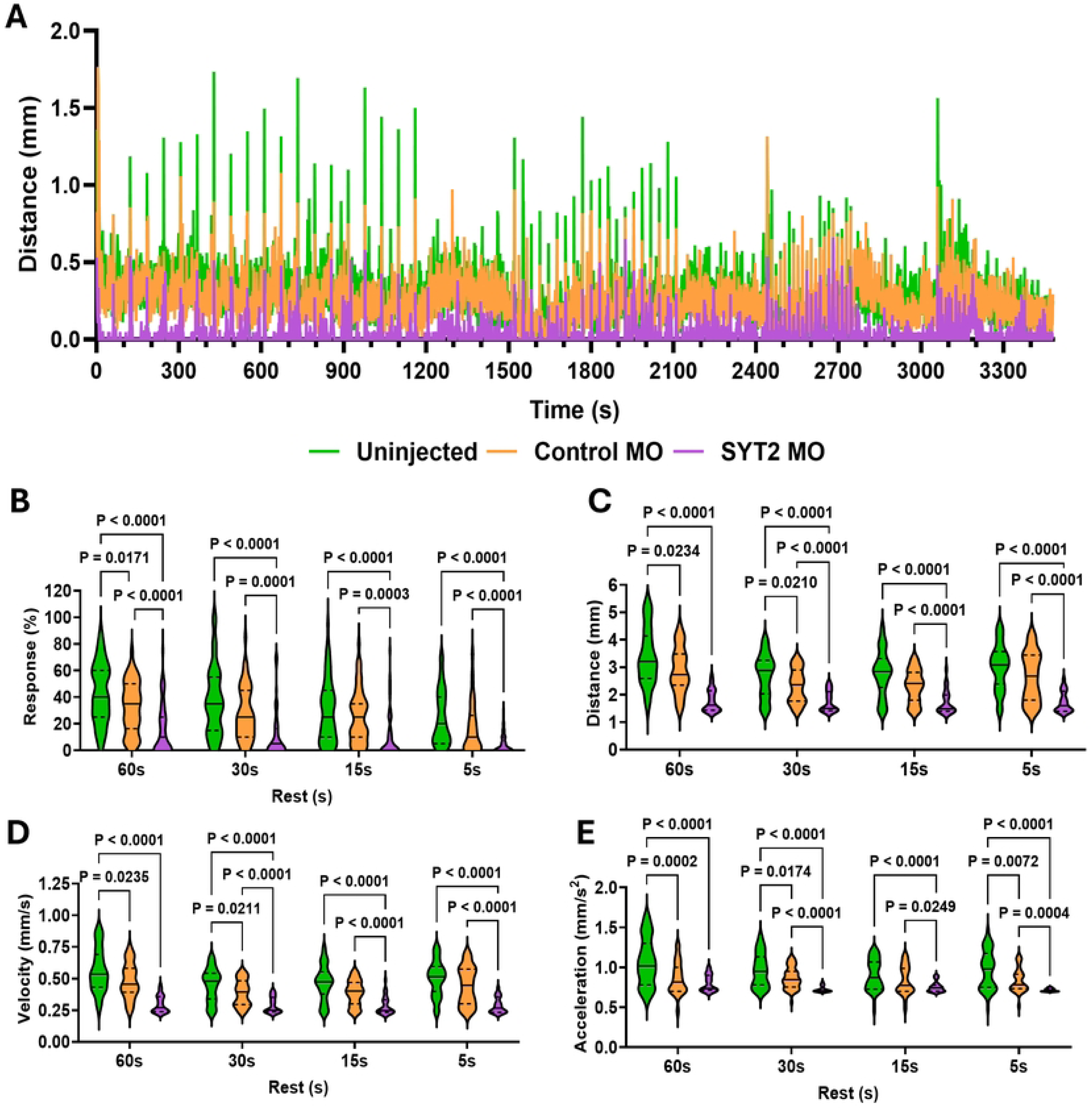
Optogenetic stimulation reveals fatigable muscle weakness in *syt2* morphants. **(A)** Trace of distance travelled during the optogenetic fatigable weakness protocol at 7 dpf where larvae were plated individually in 48-well plates (300 μL E3) and placed in a Zantiks MWP system. After 10 min dark acclimation, fish received 20, 420nm LED light pulses (1 s each) with 60 s rest intervals, followed by 5 min recovery. This stimulation block was repeated with progressively shorter rest intervals (30 s, 15 s, 5 s) to induce fatigable muscle weakness. **(B)** Average response frequency to optogenetic stimulus during each rest block for all knockdown conditions shows that *syt2* morphants respond significantly less at all rest blocks when compared to controls (uninjected, *n* = 61; Ctrl MO, *n* = 58; SYT2 MO, *n* = 51). **(C)** Average distance travelled and **(D)** Average velocity during and for the following 5 s post stimulus were found to be significantly decreased in *syt2* morphants when compared to controls for all rest blocks. **(E)** Average acceleration of responding larvae was also found to be significantly decreased in *syt2* morphants compared to controls. *Statistical analysis*, Mixed-effects analysis with Tukey’s multiple comparison’s test.

Average distance revealed a decline when comparing control groups to *syt2* morphants in all recovery blocks (**Fig. 4*C***). Slight declines were also seen between uninjected and control morphants in the 60 s and 30 s recovery blocks (**Fig. 4*C***). Differences were also observed from block to block in the uninjected and control conditions (**Fig. S2*H***). The uninjected larvae show a general decline from the 60 s recovery block, a similar pattern can be seen in the control morphants although the 15 s recovery block showed no significant difference from the 60 s recovery block (**Fig. S2*H***). As with the uninjected group, the *syt2* morphants showed significant decrease in distance travelled when comparing the 60 s recovery block to all other recovery blocks, although this effect is somewhat muted in *syt2* morphants (**Fig. S2*H***). The *syt2* morphants in the 30 s recovery block showed significantly more distance travelled than the 5 s recovery block which is likely caused by the dramatic decrease in response to stimulus in the 5 s recovery condition (**Fig. S2*H***). Average velocity found identical trends (**Fig. 4*D***) to those found in distance travelled (**Fig. 4*C***) when comparing *syt2* morphants to controls. However, looking at the block-to-block comparisons, we see significant differences in both the uninjected and control morphants when comparing the 60 s recovery block to both the 30 s and 15 s (**Fig. S2*I***). There were no differences seen in the average velocity of *syt2* morphants from block-to-block, likely due to there being little to no movement in any recovery block (**Fig. S2*I***). When looking at evoked acceleration in *syt2* morphants, we observed a sharp decrease as compared to all control clutch mates during all rest blocks (**Fig. 4*E***). Additionally, it was seen that uninjected fish had significantly higher acceleration than control morphants in the 60, 30, and 5 s recovery blocks (**Fig. 4*E***). However, block-to-block comparisons found no differences in evoked acceleration except between the 60 s and 15 s recovery blocks in uninjected larvae (**Fig. S2*J***).

### Optogenetic stimulation effectively activates motoneurons to evoke EPCs recorded from muscle fibers

We next sought to determine whether short pulses of blue light were successful in exciting motoneurons innervating the ventral fast muscle fibers selected for patch-clamp. We delivered 5-ms blue-light (480 nm) pulses over approximately one spinal segment (**Fig. S1**) and found this to be sufficient to evoke EPCs in muscle fibers (**Fig. 5**). To quantify the amplitude of evoked EPCs, we delivered five 5 ms blue-light pulses with a 52-second inter-pulse interval and averaged the amplitude of the five EPCs evoked in response to the optogenetic stimulation (**Fig. 5*A-D***). No differences in the average amplitude of evoked EPCs across groups were observed (**Fig. 5*D***; uninjected: 2,229.29 ± 1961.99 pA; control MO: 2,064.28 ± 1,473.34 pA; *syt2* MO: 1,440.52 ± 1,030.90 pA). The latency of EPCs (measured from onset of blue-light pulse to the peak of EPC) also did not differ across groups (**Fig. 5*E***; uninjected: 5.06 ± 1.30 ms; control MO: 4.80 ± 1.23 ms; *syt2* MO: 4.92 ± 1.33 ms). Finally, we utilized the evoked EPC amplitudes to estimate release properties at the NMJ (specified in the methods section). We found a significant decrease in the estimate of quantal content in *syt2* MO fish compared to both control groups (**Fig. 5*F***; uninjected: 5.30 ± 3.39; control MO: 6.08 ± 3.80; *syt2* MO: 2.66 ± 1.55). On the other hand, we observed no effects to estimates of number of release sites (**Fig. 5*G***; uninjected: 12.3 ± 10.1; control MO: 8.90 ± 6.85; *syt2* MO: 5.62 ± 4.08) or release probability (**Fig. 5*H***; uninjected: 0.642 ± 0.254; control MO: 0.745 ± 0.200; *syt2* MO: 0.662 ± 0.271). These results demonstrate defective NMJ signal transmission in *syt2* MO fish, supporting the usefulness of this optogenetic approach to estimate NMJ release properties.

**Figure 5.**
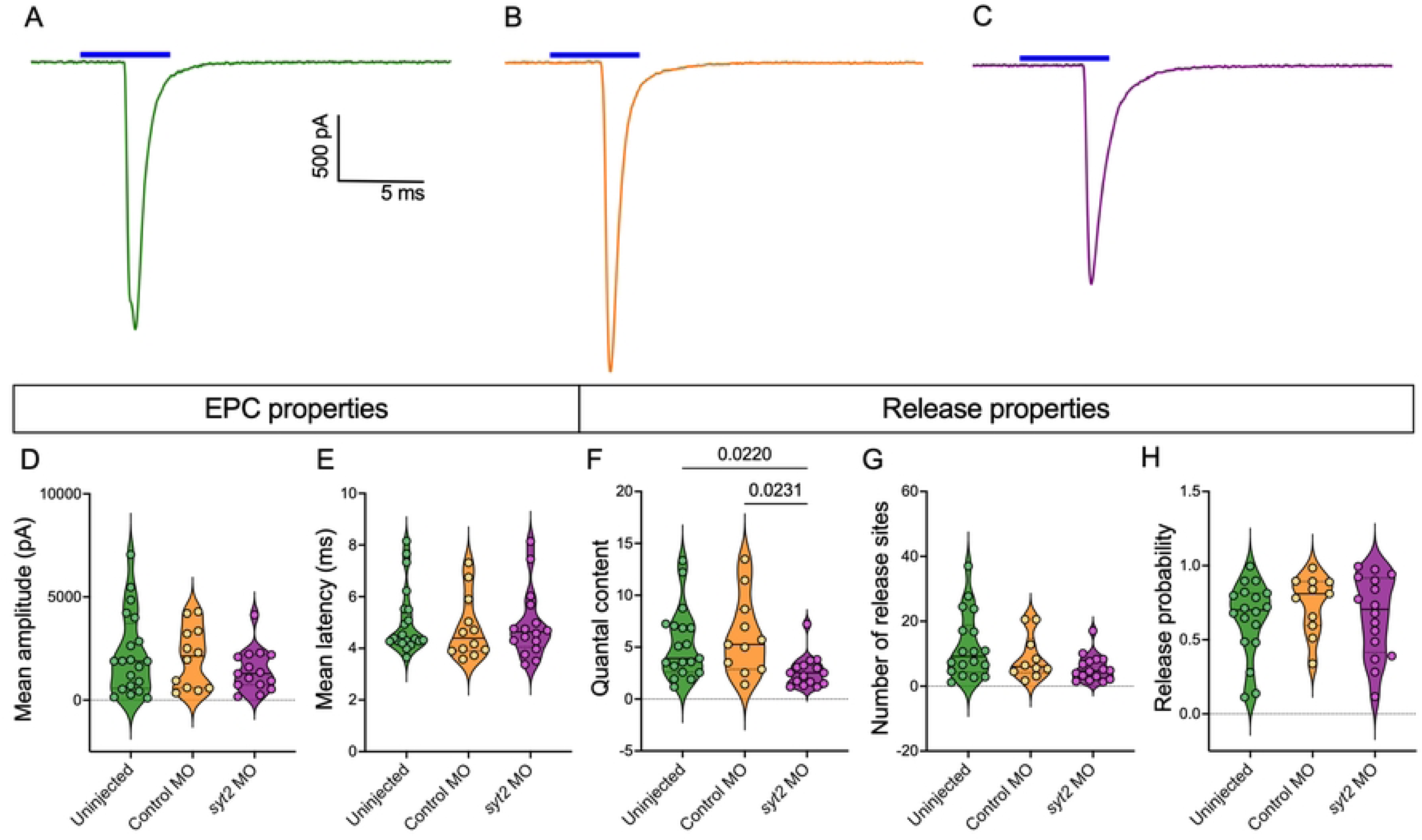
Optogenetic activation of motoneurons evokes EPCs in fast muscle fibers. **(A-C)** Representative traces of evoked end-plate currents recorded from ventral fast muscle fibers in response to a 5-ms blue light pulse in uninjected (**A**; *n* = 20 muscle fibers; *N* = 11 animals), control MO (**B**; *n* = 12; *N* = 7), and *syt2* KD (**C**; *n* = 16; *N* = 10) fish. **(D)** Average amplitude of evoked EPCs (P = 0.5648). **(E)** Average latency of EPC from onset of optogenetic stimulation (P = 0.6976; sample sizes as in ***A***-***C***). **(F)** Estimate of quantal content (uninjected versus control MO, P > 0.9999; uninjected versus *syt2* KD, P = 0.02220; control MO versus *syt2* KD, P = 0.0231; uninjected, *n* = 19, *N* = 10; control MO, *n* = 11, *N* = 7; *syt2* MO, *n* = 15, *N* = 10). **(G)** Estimated number of release sites (P = 0.0954; uninjected, *n* = 18, *N* = 10; control MO, *n* = 10, *N* = 7; otherwise, sample sizes same as in ***A***-***C***). **(H)** Estimated release probability (*P* = 0.5425; uninjected, *n* = 18, *N* = 10; control MO, *n* = 11, *N* = 7; otherwise, sample sizes same as in **A**-**C**). *Statistical analysis*, Kruskal-Wallis test with Dunn’s test for multiple comparisons (**D**-**G**); Ordinary one-way ANOVA (**H**).

### Using optogenetic stimulation to assess steady-state depression at the neuromuscular junction

We next wanted to determine whether optogenetic motoneuron stimulation could provide insight into the assessment of steady-state depression at the NMJ, as performed with electrical motoneuron stimulation (21). While Wen et al. did not assess steady-state depression in *syt2* MO fish (19), we deemed it useful to determine the feasibility of assessing steady-state depression with this optogenetic approach. Similarly to Wen et al. (2016), to assess steady-state depression, we introduced thirty 5-ms blue light pulses delivered at a frequency of 20 Hz (**Fig. S3**). We find no significant difference in the level of steady-state depression (measured as the mean amplitude of the last 25 EPCs over the amplitude of the first EPC) between groups (**Fig. S3*D***; uninjected: 0.586 ± 0.417; control MO: 0.682 ± 0.417; *syt2*MO: 0.647 ± 0.478).

Furthermore, we noticed instances when this high frequency optogenetic stimulation of motoneurons failed to result in an EPC recorded in the muscle fiber. When we quantified these failures, we found no difference across groups (**Fig. S3*E***; uninjected: 5.79 ± 5.32; control MO: 2.29 ± 3.68; *syt2* MO: 8.43 ± 6.75).

We next measured the amplitudes of the last 10 EPCs evoked during the 20 Hz stimulation, finding a difference only between control MO and *syt2* MO groups concerning the average amplitude (**Fig. S3*F***; uninjected: 1232.06 ± 908.44 pA; control MO: 526.26 ± 256.17 pA; *syt2* MO: 506.57 ± 369.55 pA). When looking into the variability of the last 10 EPC amplitudes, we observed no difference in the standard deviation of amplitudes across groups (**Fig. S3*G***; uninjected: 816.43 ± 461.50 pA; control MO: 443.22 ± 310.40 pA; *syt2* MO: 474.41 ± 356.86 pA). We did however observe a significant increase in the coefficient of variation of the *syt2* MO amplitudes compared to both control groups (**Fig. S3*H***; uninjected: 0.644 ± 0.308; control MO: 0.526 ± 0.233; *syt2* MO: 1.10 ± 0.700). These data suggest that while levels of steady-state depression at *syt2* MO NMJs are similar to controls, there is increased variability in the amplitude of EPCs at steady-state. Overall, these results demonstrate that optogenetic stimulation of motoneurons at 20 Hz in ChR2-mCherry zebrafish may be useful in assessing features of steady-state depression at the NMJ in other zebrafish models of CMS, though instances of failures must be taken into consideration.

### Using optogenetic stimulation to assess contributions of synchronous and asynchronous release at the NMJ

We next assessed contributions of synchronous and asynchronous release using our optogenetic motoneuron stimulation approach (**Fig. 6**). Synchronous neurotransmitter release induced by the membrane depolarization from motoneuron action potentials gives way to asynchronous release, which is not synchronous with incoming action potentials, during periods of high frequency activity. This asynchronous release is due to leftover increased levels of calcium in between high frequency action potentials (40). Synchronous and asynchronous release at the NMJ in *syt2* MO zebrafish has already been described using electrical stimulation of CaP motoneurons during whole-cell patch-clamp experiments (19). Wen et al. (2010) evaluated the effect of *syt2* MO on synchronous versus asynchronous release at the NMJ by electrically stimulating the CaP motoneuron at a frequency of 100 Hz for 10 s. Given the temporal constraints of our ChR2 line, we had to modify the stimulation protocol described in Wen et al. (2010) and deliver 5-ms blue-light pulses at a frequency of 20 Hz instead of 100 Hz for 10 s (**Fig. 6*A***).

**Figure 6.**
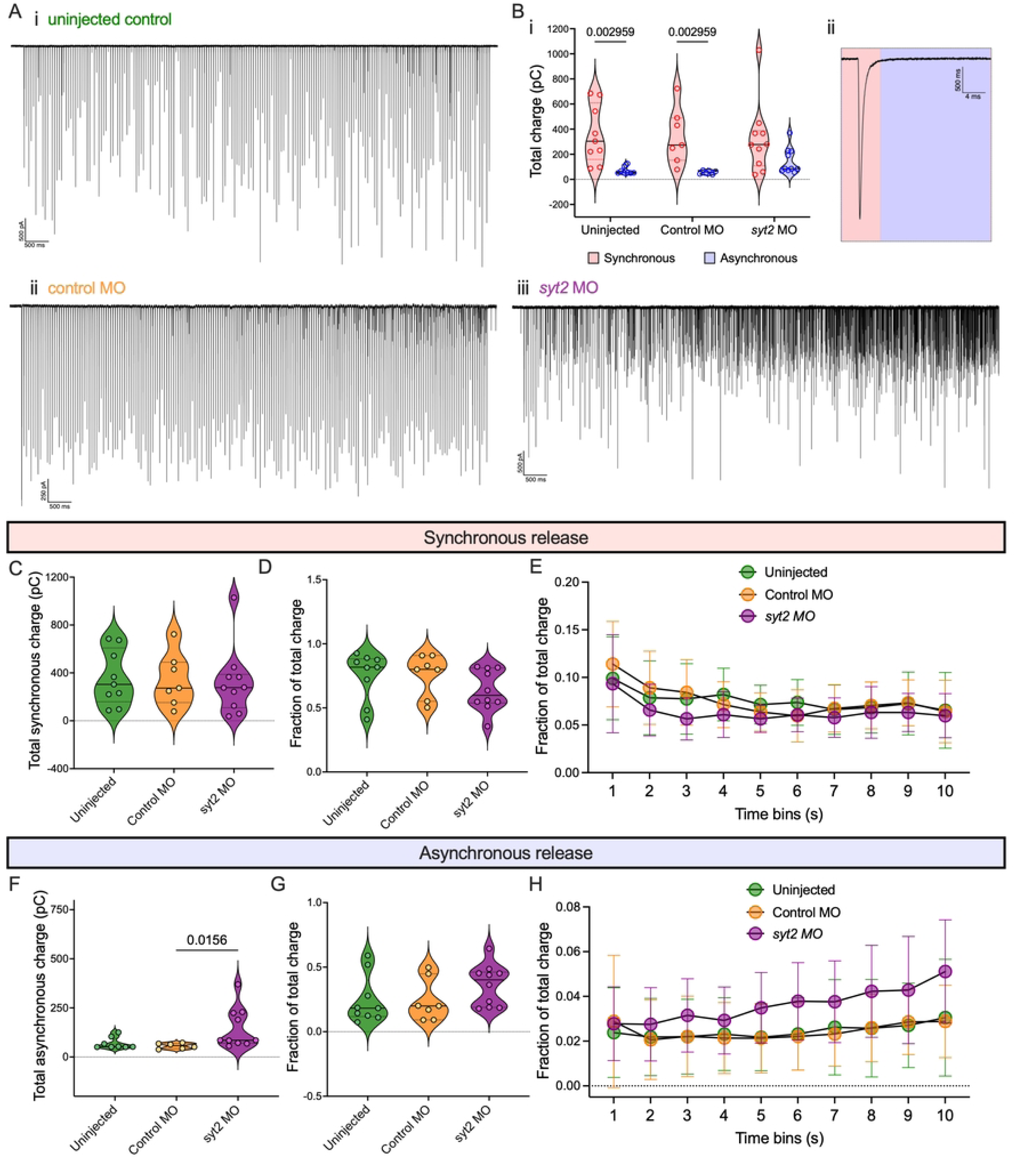
Using optogenetic stimulation to assess contributions of synchronous and asynchronous release at the NMJ. **(A)** Representative traces of EPCs recorded from ventral fast muscle fibers during 10-s 20 Hz stimulation of 5 ms blue light pulses in uninjected (**i**; *n* = 9 muscle fibers; *N* = 8 animals), control MO (**ii**; *n* = 7; *N* = 5), and *syt2* MO (**iii**; *n* = 10; *N* = 8). **(B)** Comparison of (***i***) total charge corresponding to (***ii***) synchronous (pink) and asynchronous release (blue) (uninjected, P = 0.002959; control MO, P = 0.002959; syt2 MO, P = 0.1051). **(C)** Total charge corresponding to synchronous events over the 10-s stimulation (P = 0.9149). **(D)** Fraction of total charge corresponding to synchronous events during the 10-s stimulation (P = 0.1811). **(E)** Fractional synchronous release across 1-s time bins during the 10-s stimulation (treatment group effect, P = 0.2457). **(F)** Total charge corresponding to asynchronous events over the 10-s stimulation (uninjected versus control MO, P > 0.9999; uninjected versus syt2 MO, P = 0.0533; control MO versus syt2 MO, P = 0.0156). **(G)** Fraction of total charge corresponding to asynchronous events during the 10-s stimulation (P = 0.1811). **(H)** Fractional synchronous release across 1-second time bins during the 10-s stimulation (treatment group effect: P = 0.2457). *Statistical analysis*, multiple Mann-Whitney tests with Holm-Šídák’s multiple comparisons test (**B**), Kruskal-Wallis test with Dunn’s test for multiple comparisons (**C, D, F, G**) Mixed effects analysis (**E**, **H**).

We assessed contributions of synchronous and asynchronous release by calculating the charge transfer, by performing the time integral of the current signal, at the NMJ between successive light pulses. Charge measured within the first 15 ms of the onset of the light pulse was attributed to synchronous release, whereas charge measured during the remaining 35 ms of the 50-ms interpulse interval was attributed to asynchronous release (**Fig. 6*Bii***). The total contribution of asynchronous events compared to synchronous events was lower in both control groups, but not in the *syt2* MO group (**Fig. 6*Bi***). Total synchronous (uninjected: 356.4 ± 229.3 pC; control MO: 342.3 ± 221.3 pC; *syt2* MO: 323.2 ± 282.8 pC) and asynchronous (uninjected: 69.4 ± 29.2 pC; control MO: 56.23 ± 14.0 pC; *syt2* MO: 146.4 ± 103.3 pC) release were comparable across groups (**Fig. 6*C,F***). We next compared the contribution of synchronous and asynchronous events to total release. The fraction of synchronous release over total release did not differ between groups (**Fig. 6*D***; uninjected: 0.755 ± 0.189; control MO: 0.757 ± 0.164; *syt2* MO: 0.637 ± 0.159). Similarly, no differences in total fractional asynchronous release were observed between groups (**Fig. 6*G***; uninjected: 0.245 ± 0.186; control MO: 0.243 ± 0.164; *syt2* MO: 0.363 ± 0.159).

Finally, we grouped synchronous and asynchronous charge measurements into ten 1-s bins corresponding to the 10-second stimulation protocol to gain a sense of their contributions over time. While stimulation time had a significant decreasing effect on fractional charge attributed to synchronous release, no differences in fractional synchronous release over time were observed across groups (**Fig. 6*E***). This was also the case for fractional asynchronous release where stimulation time had an overall increasing effect, but no differences were observed across groups (**Fig. 6*H***). Overall, these data demonstrate the optogenetic approach’s ability to assess differences in synchronous and asynchronous release at the NMJ in *syt2* MO fish, albeit with some limitations to consider.

### SNARE Complex and other CMS proteins are downregulated in syt2 morphants

Although there are 40 genes connected to CMS, the response of CMS and other NMJ-specific genes to knockdown of any one CMS gene is not fully understood. Examining the expression of these genes in the context of this *syt2* morpholino model could reveal hidden pathomechanisms and novel therapeutic targets for the treatment of presynaptic CMS. To examine NMJ gene expression in *syt2* morphants, we developed a NanoString® panel to quantify transcripts of NMJ-relevant genes. We assembled target genes by identifying two transcriptomic studies that isolated NMJ-specific genes (36,37) and by generating protein–protein interaction networks from the 35 known CMS genes (*AGRN*, *ALG14*, *ALG2*, *CHAT*, *CHD8*, *CHRNA1*, *CHRNB1*, *CHRND*, *CHRNE*, *CHRNG*, *COL13A1*, *COLQ*, *DOK7*, *DPAGT1*, *GFPT1*, *GMPPB*, *LAMA5*, *LAMB2*, *LRP4*, *MUSK*, *MYO9A*, *PLEC*, *PREPL*, *PURA*, *RAPSN*, *RPH3A*, *SCN4A*, *SLC18A3*, *SLC25A1*, *SLC5A7*, *SNAP25*, *SYT2*, *TOR1AIP1*, *UNC13A*, *VAMP1*, *MACF1*) using STRING and IPA software (1,41,42). These networks were expanded to include the closest ∼200 NMJ and CMS related genes to reveal pathways not previously linked to NMJ function. In addition, to help improve signal of NMJ relevant genes conserved across species, only genes present in at least two species-derived networks (*Homo sapiens*, *Mus musculus*, *Rattus norvegicus*, *Danio rerio*) which had zebrafish orthologues were included in the final panel (**Fig. 7*A***; full target and probe lists provided in the **Supplementary Material**). We applied this panel to whole larval RNA extracts from 3 dpf ChR2 larvae across all experimental groups (uninjected, control MO, *syt2* morphants). Control MO injection did not significantly alter expression relative to uninjected larvae, therefore these datasets were combined and used as a single reference dataset when analyzing *syt2* morphant gene expression (**Fig. S4*A***). Separate comparisons of each data set revealed similar DEGs (**Fig. S4*B*-*C***). Using the combined uninjected + control MO dataset as a reference, we identified several significantly downregulated genes in *syt2* morphants (**Fig. 7*B***). Downregulated genes did not include *syt2* as samples were collected from morphants injected with translation-blocking mo. Downregulated genes included *sypb, calm1a, myo9a, vamp2, snap25b, stxbp1b, unc13a,* and *slc17a7a*. Of these genes *myo9ab*, *snap25b*, and *unc13a* are known CMS genes and *snap25b* and *unc13a* cause LEMS-Like CMS (43,44). These genes all code for various synaptic vesicle release machinery. *SNAP25B* is an autosomal dominant gene that codes for a core Q-SNARE protein that anchors and primes synaptic vesicles to the pre-synaptic membrane (44,45). *UNC13A* codes for the syntaxin stabilizing protein Munc13-1 (43,45). Lastly, *MYO9A* is a CMS gene that codes for myosin IXA, a key protein in NMJ development and the excretion of the agrin signalling molecule (23,26). Downregulated genes not directly linked to CMS included *sypb*, *caml1a*, *vamp2*, *stxbp1*, and *slc17a7*. Mutations in several of these genes are known to cause human disease according to the OMIM database.

**Figure 7.**
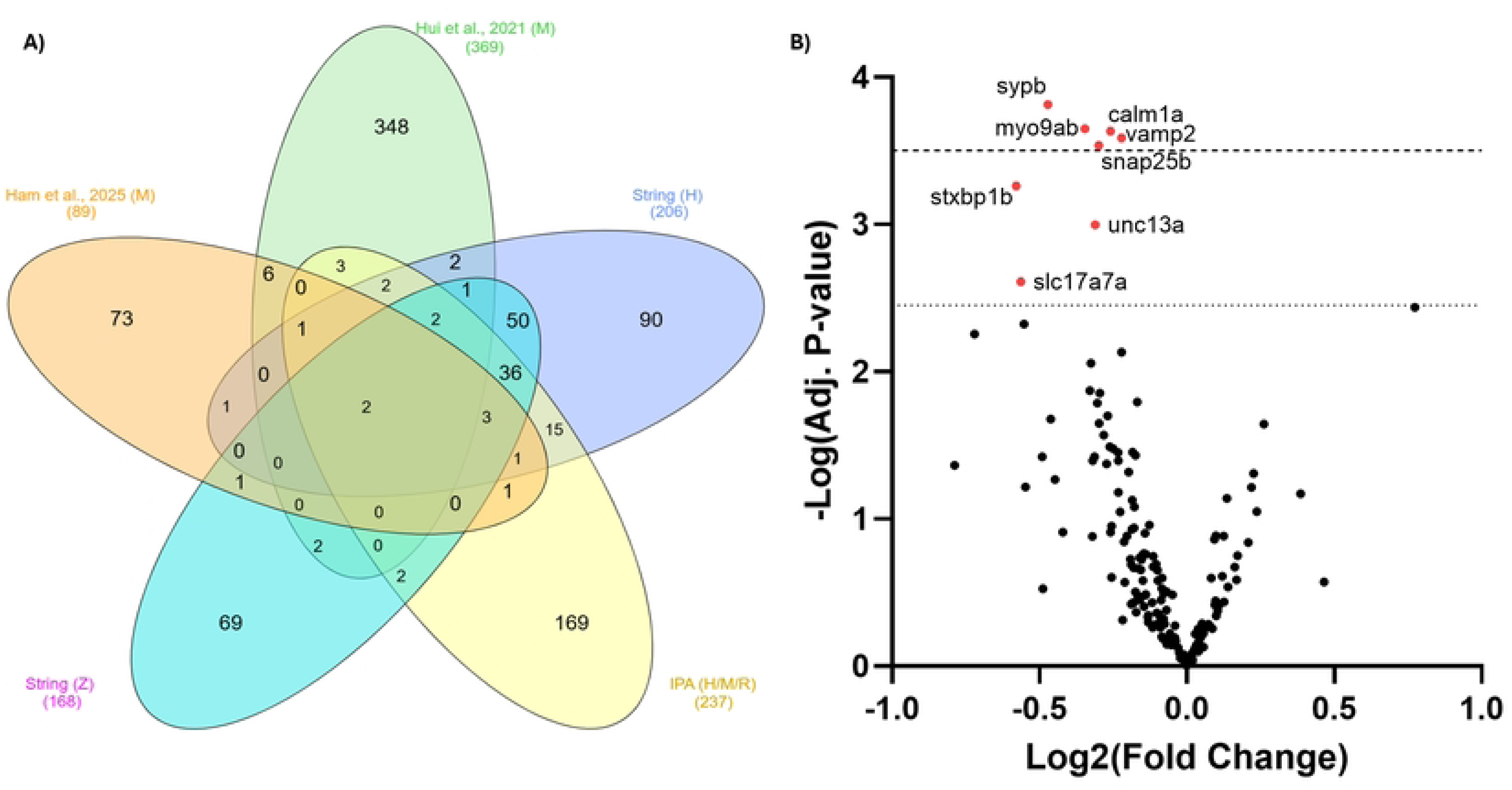
Dysregulation of active zone proteins alter neuromuscular junction signaling dynamics in syt2 morphants. **(A)** Venn diagram of common genes from protein-protein interaction networks generated with STRING, IPA, or from neuromuscular junction-specific RNA-seq datasets from Human (H), mouse (M), rat (R), and zebrafish (Z) found in literature. The expression panel developed from these sets of data included only genes found in at least 2 lists and had to have zebrafish orthologs. **(B)** Volcano plot visualizing differentially expressed genes (DEGs) in *syt2* morphants (red = decreased expression. Dotted line, *adj*. P = 0.50; Dashed line, *adj*. P = 0.10).

Firstly, the human ortholog for *sypb* is *SYP* and codes for synaptophysin, a protein which helps to ensure efficient synaptic vesicle endocytosis, is associated with intellectual developmental disorder and X-linked 96 (OMIM:313475). Next, *calm1a*, which codes for calmodulin, is a cytoplasmic calcium sensor that participates in many Ca2+-dependent processes in motor neurons. Zebrafish *stxbp1b* is one of the orthologues of *STXBP1*. This gene codes for Munc18-1 an important regulatory component of SNARE complex proteins (46–48). VAMP2 is a core SNARE protein that is essential for fast, Ca2+-dependent release at synapses and is modulated by synaptophysin to help position it for efficient neurotransmitter release in neurons located in the central nervous system (49).

## Discussion

In this study, we generated *syt2* MO knockdowns in ChR2-mCherry (32,33) fish to investigate a multilevel, optogenetic approach for the assessment of locomotor and NMJ function in the context of screening novel CMS gene candidates. We chose to investigate *syt2* as it is a known CMS gene and many of its locomotor, electrophysiological, and genetic effects have been characterized in both mice and zebrafish (19,50–56). We sought to better define muscle fatigability in CMS, relate behavioral and electrophysiological data through a widespread optogenetic stimulation of motor neurons, and to further characterize the NMJ-related genetic landscape to further postulate novel pathomechanisms, identify novel gene candidates, and find novel pathways to target therapeutically for CMS treatment.

### Interpretation of behaviour and endplate dynamics upon optogenetic stimulation

A key feature of our approach is that similar stimulus is used to elicit motor responses at both behavioural and synaptic levels. This allows us to better relate specific electrophysiological deficits to behavioural ones. Defects to NMJ function in *syt2* knockdown fish were first revealed by investigating the NMJ of caudal primary (CaP) motoneurons to fast ventral muscle fibers (19). In contrast to that previous study, our optogenetic stimulation is not limited to the CaP motoneuron. It is important to note that while restriction of optogenetic excitation to individual motoneurons can be performed with advanced microscopy techniques, during both behavioral and electrophysiological studies, we illuminated either the entire larvae or whole spinal segments. Recorded endplate currents in the ventral fast muscle fibers are therefore likely to include signal transmission from the CaP motoneuron as well as secondary motoneurons that innervate this ventral-most quadrant of musculature due to the polyneuronal innervation of individual muscle fibers found in zebrafish (28,57,58). Indeed, the ChR2-mCherry line may express ChR2 mostly in secondary motoneurons (28), though we have observed frequent expression in primary motoneurons. Our data therefore may more accurately represent population level EPCs rather than purely CaP-associated EPCs such as those investigated previously (19,28). Furthermore, our full-segment blue light excitation approach inevitably excites any spinal interneurons expressing ChR2 as well. Indeed, we notice instances of doublet and triplet EPCs generated in response to our optogenetic stimulation (data not shown). These are likely a result of polyneuronal motoneuron activation while the first EPC generated with short latency could be a summation of multiple EPCs from more than one motoneuron. Additionally, polyneuronal activation may explain why we observed a significant decrease in response to stimulation by *syt2* morphants during the fatigable weakness assay yet the failure rate of EPCs, although highly variable, showed no differences from controls. It is possible polyneuronal activation may have an additive effect where the summation of multiple smaller signals is enough to elicit a response in the form of EPCs that would still be insufficient to elicit contraction. Also, the transient/mosaic nature of morpholino knockdown may vary the knockdown level amongst motoneurons. However, the ability to stimulate multiple neurons simultaneously may produce a more relatable readout of electrophysiological phenotype in relation to behavioral results because it encapsulates the poly-innervated nature of the zebrafish neuromuscular system. Regardless of the nature of the EPCs (whether single motoneuron driven or population driven), this approach effectively stimulates motoneurons for assessment of NMJ function at both the behavioral and synaptic level, though it is imperative that interpretations of the results take into consideration these factors.

While our measures of estimated number of release sites fell within the normal range for controls (21), our observed decreases in quantal content and endplate potential (EPP) amplitude in *syt2* morphants is in line with mouse *Syt2* knock out studies (50). Interestingly, although *syt2* is a calcium sensor where its dysfunction would be thought to cause a decrease in release probability, it remained unchanged in *syt2* morphants. In fact, CMS patients with *UNC13A* mutations, which we show has decreased expression in *syt2* morphants, also show decreased quantal content but no differences in synaptic vesicle release probability in humans (43). It is possible that decreased expression of *unc13a* explains this characteristic of *syt2* morphant phenotype and may provide possible mechanistic insight into *SYT2*-CMS.

Motoneuron population driven EPCs likely lead to increased variability in the amplitude of recorded EPCs. There is also the possibility that our optogenetic approach mainly activated secondary motoneuron at the exclusion of CaP motoneurons. As mentioned above, ChR2-mCherry fish may exhibit ChR2 expression predominantly in secondary motoneurons (28). The lower quantal content released from secondary motoneurons when compared to primary motoneurons (28) could explain our own lower estimates of these measures of neuromuscular transmission.

While we found visual evidence that CaP motoneurons express ChR2 through mCherry expression, it is nonetheless possible that optogenetic stimulation proves ineffective in consistently depolarizing the larger primary motoneuron to threshold (59). Whichever the level of optogenetic CaP motoneuron activation, this level may be consistent enough across treatment groups that comparisons to controls are still able to reveal differences in properties of population-level signal transmission.

### Temporal constraints linked to channelrhodopsin kinetics

Channelrhodopsin proteins are associated with known temporal constraints given the kinetic properties of opsins (60). High frequency stimulation protocols are important to the study of signal transmission, shedding light onto features of synaptic release, facilitation, and depression using common protocols that include the paired-pulse ratio as well as high frequency stimulation trains. While the effects of *syt2* knockdown on steady-state depression were not assessed previously (19), we chose to do so to elucidate the use of optogenetic stimulation at high frequencies for NMJ function assessment. Previous work successfully stimulated the CaP motoneuron at 20, 50, and 100 Hz using electrical stimulation to assess steady-state depression occurring at the NMJ (21). We were unable to stimulate motoneurons optogenetically at 50 Hz nor 100 Hz (data not shown) but had success with 20 Hz. This agrees with known temporal constraints of ChR2 proteins permitting reliable high frequency stimulation up to 30 Hz (Boyden *et al*., 2005). Interestingly, levels of steady-state depression observed in controls in response to optogenetic motoneuron stimulation are lower compared to electrical stimulation delivered at 20 Hz to CaP motoneurons (21). This confounding result speaks to the likelihood of secondary motoneuron activation alone or in conjunction with CaP motoneuron within a spinal segment. Neurotransmitter release from sMNs shows facilitation in response to repetitive stimulation as opposed to synaptic fatigue exhibited by primary motoneurons (28).

When we investigated the role of *syt2* in synchronous versus asynchronous release at the NMJ, our protocol had to be fundamentally different from previous investigations (19). Those electric stimulation protocols consisted of a 100 Hz train lasting 10 seconds (19) Direct comparisons to our optogenetic approach are not possible as optogenetic stimulation at 100 Hz cannot be achieved. We chose instead to deliver a 20 Hz train of 5-ms blue light pulses over the course of 10 seconds. While we did indeed see the apparition of asynchronous events towards the end of the protocol, it is unclear whether 20 Hz for this length of time is sufficient to fully capture asynchronous versus synchronous contributions. Previous work reveals that during 10-s 100 Hz stimulation, synchronous release rapidly declines as asynchronous release rapidly increases within the first few seconds of stimulation (19). Contributions from synchronous release continue to decline and those from asynchronous release begin declining mid-way through the stimulation protocol (19). With our 10-s 20 Hz optogenetic stimulation protocol, we observe a steady yet modest decrease in contributions from synchronous release with a similar steady yet modest increase in asynchronous events throughout the stimulation protocol. Lower frequency stimulation is likely to be less effective at depleting vesicle stores, making it harder to investigate effects to synchronous release. The lower frequency of our stimulation protocol is also likely to be behind the failure to notice a sharp increase in asynchronous events early within the stimulation window – 20 Hz stimulation may be ineffective in prolonging the inter-spike raises in intracellular calcium levels that trigger asynchronous release. It is possible that lower train rates may require longer stimulation durations that have not been tested in this study. Furthermore, while the CaP motoneuron exhibits a high probability of release, that of secondary motoneurons is significantly lower (28) meaning their synchronous versus asynchronous release profiles over the same time course of stimulation would not resemble the profile observed for the primary CaP motoneuron alone.

### Advantages associated with optogenetically assessing NMJ function

Traditional studies of NMJ function by electrical stimulation of individual motoneurons and recording evoked EPCs in innervated muscle fibers provide important insights into individual neuromuscular connections. While single synapse studies are imperative to the understanding of individual protein contributions to signal transmission across the NMJ, our population level approach for NMJ function assessment is more representative of the neuromuscular system in action during movement production, especially considering that motoneurons are polyneuronally innervated in zebrafish (28,57,58). The optogenetic approach employed here serves as a valuable complement to traditional NMJ studies by investigating a physiologically relevant link between behaviour and NMJ function through population-evoked EPCs.

This approach also offers flexibility to the assessment of NMJ function. While we chose to stimulate motoneurons within one segment of the spinal cord, one could choose to stimulate over more spinal segments or restrict excitation to individual motoneurons. Furthermore, the generation of fast genetically encoded calcium indicators restricted to muscle fibers could be utilized in conjunction with optogenetic activation of motoneurons like that described in this study for an increasingly less technically challenging method for NMJ function assessment, albeit bearing its own set of limitations.

### NMJ gene expression in *syt2* morphants reveals connections to synaptic recycling dynamics and activity-based phenotype

We utilized a gene-panel to look specifically at genes associated with the NMJ. Knockdown of *syt2* was accompanied by the downregulation of multiple genes essential to presynaptic vesicle cycling, including priming and SNARE machinery (*unc13a, stxbp1, snap25b, vamp2,* and *sypb*), Ca2+-dependent vesicle priming & replenishing pathways (*calm1a*), synaptic vesicle trafficking (*myo9ab*), and neurotransmitter loading (*slc17a7a*). Overall, this transcriptional profile is consistent with the phenotype characteristics seen in our behavioral and electrophysiological observations of reduced optogenetic response rate, reduced quantal content, decreased evoked EPC amplitude, and increased asynchronous release.

Among these DEGs, calmodulin (*calm1a*) is of particular interest as it has many binding partners and a known inverse regulatory relationship with *syt2* in cortical neurons and multiple areas of the mouse brain (61). Binding sites for Ca2+/calmodulin have been identified in the CMS gene *MYO9A*, where it negatively regulates its motility and ATPase activity (62). Additionally, calmodulin interacts with the CMS protein MUNC-13 (*unc13a*) to regulate vesicle priming and influence short-term synaptic plasticity (63). Furthermore, deletion/downregulation of *UNC13A* results in inefficient SNARE complex formation around voltage-gated Ca2+ channels, decreases the number of readily releasable vesicles at the presynaptic terminal and effectively blocks synaptic vesicle fusion (46–48). As *syt2* is not a transcription factor, it is not surprising that expression differences were limited to genes that have a direct role in, or regulation of, synaptic transmission. However, studies on *Syt2* knockout mice showed different results when measuring protein levels as compared to transcript levels, namely of Snap25, Syt-1, and synaptobrevin (also known as Syb or Vamp2) (50). We found that whole-larva transcript expression levels of both *vamp2* and *snap25b* were significantly downregulated in *syt2* morphants as compared to *Syt2* knockout mice who had unchanged protein levels in CNS tissues (50). This highlights potential tissue-specific differences or genetic compensation effects due to genome replication events in zebrafish (64). Considering these differences, dysregulated genes in the present study may be representative of peripheral tissues and highlight their function in the neuromuscular system in the context of *syt2* knockdown. In all, these DEGs suggest compounding defects in all parts of the vesicle cycle led to a CMS phenotype rather than an isolated failure of Ca2+-sensing by *syt2*.

### Concluding remarks

The optogenetic approach described in this study to excite motoneurons using the ChR2-mCherry line effectively reveals important differences in *syt2* morphants described previously, though with some discrepancies due to both the nature of optogenetics as well as our method of stimulation. Keeping these limitations in my mind, our results support our approach as a potentially useful, relatively facile, and high throughput means to screen for gene candidates underlying genetically undiagnosed cases of CMS.

While we found that optogenetic motoneuron activation has its limitations, our genetic analysis revealed interesting results that require further investigation into the compounding potential defects that lead to pathogenesis in *SYT2*-CMS. Our multilevel platform may prove to be a useful tool for the rapid screening of novel CMS genes, a process that typically requires extensive behavioral, electrophysiological and genetic techniques.

## Acknowledgements

The SV2 and ZNP-1 monoclonal antibodies were developed by the Harvard Medical School and the University of Oregon respectively, was obtained from the Developmental Studies Hybridoma Bank, created by the NICHD of the NIH and maintained at The University of Iowa, Department of Biology, Iowa City, IA 52242. The 1020:Gal4;UAS:ChR2(H134)-mCherry was provided by Dr. Tod Thiele.

HL receives support from the Canadian Institutes of Health Research (CIHR) for Foundation Grant FDN-167281 (Precision Health for Neuromuscular Diseases), Transnational Team Grant ERT-174211 (ProDGNE) and Network Grant OR2-189333 (NMD4C), from the Canada Foundation for Innovation (CFI-JELF 38412), the Canada Research Chairs program (Canada Research Chair in Neuromuscular Genomics and Health, 950-232279), the European Commission (Grant # 101080249) and the Canada Research Coordinating Committee New Frontiers in Research Fund (NFRFG-2022-00033) for SIMPATHIC, and from the Government of Canada Canada First Research Excellence Fund (CFREF) for the Brain-Heart Interconnectome (CFREF-2022-00007).

TVB and SFG receive support from the Natural Sciences and Engineering Research Council of Canada (NSERC): NSERC Discovery Grant, Grant Number: RGPIN-2022-03898 (to TVB); NSERC Canadian Graduate Scholarship M award, Award Number: NSERC 553401-2020 (to SFG); NSERC Postgraduate Scholarship D, Award Number: 569969-2022 (to SFG).

## Supplementary Materials

**Figure S1. Channelrhodopsin expression restricted to spinal interneurons and motoneurons in 1020:Gal4;UAS:ChR2(H134)-mCherry fish. (A)** Schematic oversimplification of experimental setup wherein 480 nm optogenetic excitation is delivered to an entire spinal segment within which reside the motoneurons innervating the patch-clamped fast muscle fiber. **(B-D)** Example images taken from 3 individual transgenic zebrafish to show variability of channelrhodopsin (mCherry) expression. **(B)** Image of the spinal cord containing mCherry-positive spinal neurons in a 2 dpf zebrafish. **(C)** Image of the spinal cord containing mCherry-positive spinal neurons and dorsal-most section of the ventral musculature where motoneuron axons are present in a 4 dpf zebrafish. **(D)** Image of the spinal cord containing mCherry-positive spinal neurons and dorsal-most section of the ventral musculature in a different 4 dpf zebrafish from **C**. Boxed area (*i*) is enlarged in (*ii*). Arrow heads are pointing to ventral roots containing motoneuron axons that exist the spinal cord to innervate muscles. MN: motoneuron. Image created using biorender.com.

**Figure S2. Further behavioral characterization of *syt2* morphants. (A)** representative photos of *syt2* morphants on a WT background. Scale bar: 500 µm. **(B)** Measurements of the proportion of AChR clusters which colocalize with ZNP-1 (uninjected = 0.459 ± 0.0170 ; control MO = 0.636 ± 0.0811; *syt2* MO = 0.0588 ± 0.0106; 2-way ANNOVA) and **(C)** shows the proportion of ZNP-1 spots with colocalizing AChRs ZNP-1 (uninjected = 0.563 ± 0.119 ; control MO = 0.595 ± 0.0381; *syt2* MO = 0.134 ± 0.0886; 2-way ANNOVA). Additionally, a touch-response assay was conducted by prodding 3 dpf larvae on the back of the head and recording the **(D)** distance they moved until stopping (uninjected = 100 ± 48.9 %; control MO = 82.4 ± 32.1 %; *syt2* MO = 57.1 ± 30.7 %), and **(E)** their initial acceleration away from the physical stimulus (uninjected = 100 ± 37.8 % ; control MO = 115 ± 45.5 %); *syt2* MO = 117 ± 43.6 %). **(F)** Total distance travelled, normalized to uninjected controls during the light phase of the light-dark transition assay (uninjected = 100 ± 123 %; control MO = 26.2 ± 42.0 %; *syt2* MO = 12.0 ± 22.9 %). **G-J)** Shows analysis of the response rate **(G)**, distance travelled after stimulus **(H)**, average velocity after stimulus **(I)**, and initial acceleration after stimulus of responding larvae during the fatigable weakness assay grouped by recovery block within each phenotype. All measurements are shown as: mean ± SD.

**Figure S3. Using optogenetic stimulation to estimate steady-state depression at the NMJ. *A-C***, Representative traces of EPCs recorded from ventral fast muscle fibers elicited in response to thirty 5-ms pulses delivered at 20 Hz in uninjected **(A)**, control MO **(B)**, and *syt2* MO fish **(C)**. **(D)** Measure of steady-state depression (P = 0.8586; uninjected, *n* = 15 muscle fibers; *N* = 10 animals; control MO, *n* = 8; *N* = 5; *syt2* MO, *n* = 16; *N* = 10). **(E)** Number of failures (i.e. no EPC) during stimulation train (P = 0.0800; uninjected, *n* = 15; *N* = 10; control MO, *n* = 8; *N* = 5; *syt2* MO, *n* = 17; *N* = 10). **(F)** Average amplitude of the last 10 EPCs (uninjected versus syt2 MO, P = 0.0397; uninjected versus control MO, P = 0.2259; control MO versus syt2 MO, P > 0.9999; control MO, *n* = 7, *N* = 4; otherwise, sample sizes same as in **D**). **(G)** Standard deviation of the amplitudes of the last 10 EPCs (P = 0.0339; uninjected versus syt2 MO, P = 0.0528; uninjected versus control MO, P = 0.0906; control MO versus syt2 MO: P = 0.9817; sample sizes same as in **(D)**. ***H***, Coefficient of variation of the amplitudes of the last 10 EPCs (uninjected versus syt2 MO, P = 0.0439; uninjected versus control MO, P = 0.8629; control MO versus syt2 MO, P = 0.0428; sample sizes same as in **D**). *Statistical analysis,* One-way ANOVA with Tukey’s multiple comparisons test **(G**, **H)** and Kruskal-Wallis test with Dunn’s test for multiple comparisons **(D**-**F)**.

**Figure S4. Dysregulated genes in *syt2* morphant zebrafish**. **(A)** volcano plot showing no significant differences in NMJ-gene expression between uninjected and control injected experimental conditions showing that these datasets may be combined as one common reference to compare with *syt2* morphant expression data. **(B-C)** Show differentially expressed genes (DEGs) when comparing *syt2* morphants (syt2 MO) to **(B)** uninjected larvae and **(C)** control injected (Ctrl MO) larvae. **(D)** Individual bar charts representing the Log2 fold change in expression of each DEG found in the main results section.

## Notes

### Competing Interest Statement

The authors have declared no competing interest.

## References

1. Ohno K, Ohkawara B, Shen XM, Selcen D, Engel AG. Clinical and Pathologic Features of Congenital Myasthenic Syndromes Caused by 35 Genes—A Comprehensive Review. International Journal of Molecular Sciences 2023, Vol 24, Page 3730. 2023 Feb 13;24(4):3730. doi:10.3390/IJMS24043730

2. Ohno K, Ito M, Ohkawara B. Review of 40 genes causing congenital myasthenic syndromes. J Hum Genet. 2025 Jun 18;1–10. doi:10.1038/S10038-025-01355-9;TECHMETA=23,45;SUBJMETA=2056,308,374,375,692,699;KWRD=GENETICS+RESEARCH,NEUROMUSCULAR+DISEASE

3. Ramdas S, Beeson D, Dong YY. Congenital myasthenic syndromes: increasingly complex. Curr Opin Neurol. 2024 Oct 1;37(5):493–501. doi:10.1097/WCO.0000000000001300 PubMed PMID: 39051439.

4. Abicht A, Müller J, Lochmüller H. Congenital Myasthenic Syndromes. GeneReviews® [Internet]. 1993 [cited 2019 Oct 8]. Available from: http://www.ncbi.nlm.nih.gov/pubmed/20301347 PubMed PMID: 20301347.

5. Engel AG, Shen XM, Selcen D, Sine SM. Congenital myasthenic syndromes: pathogenesis, diagnosis, and treatment. Lancet Neurol. 2015 Apr 1;14(4):420–34. doi:10.1016/S1474-4422(14)70201-7

6. Finlayson S, Beeson D, Palace J. Congenital myasthenic syndromes: an update. Pract Neurol. 2013 Apr 1;13(2):80–91. doi:10.1136/PRACTNEUROL-2012-000404 PubMed PMID: 23468559.

7. Nasevicius A, Ekker SC. Effective targeted gene ‘knockdown’ in zebrafish. Nature Genetics 2000 26:2. 2000 Oct;26(2):216–20. doi:10.1038/79951 PubMed PMID: 11017081.

8. Chang N, Sun C, Gao L, Zhu D, Xu X, Zhu X, et al. Genome editing with RNA-guided Cas9 nuclease in Zebrafish embryos. Cell Research 2013 23:4. 2013 Mar 26;23(4):465–72. doi:10.1038/cr.2013.45 PubMed PMID: 23528705.

9. Hwang WY, Fu Y, Reyon D, Maeder ML, Tsai SQ, Sander JD, et al. Efficient genome editing in zebrafish using a CRISPR-Cas system. Nature Biotechnology 2013 31:3. 2013 Jan 29;31(3):227–9. doi:10.1038/nbt.2501 PubMed PMID: 23360964.

10. Jao LE, Wente SR, Chen W. Efficient multiplex biallelic zebrafish genome editing using a CRISPR nuclease system. Proc Natl Acad Sci U S A. 2013 Aug 20;110(34):13904–9. doi:10.1073/PNAS.1308335110;WGROUP:STRING:PUBLICATION PubMed PMID: 23918387.

11. Kroll F, Powell GT, Ghosh M, Gestri G, Antinucci P, Hearn TJ, et al. A simple and effective f0 knockout method for rapid screening of behaviour and other complex phenotypes. Elife. 2021;10:1–34. doi:10.7554/eLife.59683 PubMed PMID: 33416493.

12. Saint-Amant L, Drapeau P. Time course of the development of motor behaviors in the zebrafish embryo. J Neurobiol. 1998 Dec;37(4):622–32. doi:10.1002/(SICI)1097-4695(199812)37:4<622::AID-NEU10>3.0.CO;2-S PubMed PMID: 9858263.

13. Karuppasamy M, English KG, Henry CA, Manzini MC, Parant JM, Wright MA, et al. Standardization of zebrafish drug testing parameters for muscle diseases. Dis Model Mech. 2024 Jan 1;17(1). doi:10.1242/DMM.050339/342428 PubMed PMID: 38235578.

14. Singh J, Patten SA. Modeling neuromuscular diseases in zebrafish. Front Mol Neurosci. 2022 Dec 13;15:1054573. doi:10.3389/FNMOL.2022.1054573/XML

15. Lescouzères L, Bordignon B, Bomont P. Development of a high-throughput tailored imaging method in zebrafish to understand and treat neuromuscular diseases. Front Mol Neurosci. 2022 Sep 20;15:956582. doi:10.3389/FNMOL.2022.956582/XML/NLM

16. Berger J, Sztal T, Currie PD. Quantification of birefringence readily measures the level of muscle damage in zebrafish. Biochem Biophys Res Commun. 2012 Jul 13;423(4):785–8. doi:10.1016/J.BBRC.2012.06.040 PubMed PMID: 22713473.

17. Oliveira NAS, Pinho BR, Oliveira JMA. Swimming against ALS: How to model disease in zebrafish for pathophysiological and behavioral studies. Neurosci Biobehav Rev. 2023 May 1;148:105138. doi:10.1016/J.NEUBIOREV.2023.105138 PubMed PMID: 36933816.

18. Chia K, Klingseisen A, Sieger D, Priller J. Zebrafish as a model organism for neurodegenerative disease. Front Mol Neurosci. 2022 Oct 13;15:940484. doi:10.3389/FNMOL.2022.940484/FULL

19. Wen H, Linhoff MW, McGinley MJ, Li GL, Corson GM, Mandel G, et al. Distinct roles for two synaptotagmin isoforms in synchronous and asynchronous transmitter release at zebrafish neuromuscular junction. Proc Natl Acad Sci U S A. 2010 Aug 3;107(31):13906–11. doi:10.1073/PNAS.1008598107 PubMed PMID: 20643933.

20. Walogorsky M, Mongeon R, Wen H, Mandel G, Brehm P. Acetylcholine Receptor Gating in a Zebrafish Model for Slow-Channel Syndrome. Journal of Neuroscience. 2012 Jun 6;32(23):7941–8. doi:10.1523/JNEUROSCI.0158-12.2012 PubMed PMID: 22674269.

21. Wen H, Hubbard JM, Wang WC, Brehm P. Fatigue in rapsyn-deficient zebrafish reflects defective transmitter release. Journal of Neuroscience. 2016 Oct 19;36(42):10870–82. doi:10.1523/JNEUROSCI.0505-16.2016 PubMed PMID: 27798141.

22. McMacken GM, Spendiff S, Whittaker RG, O’Connor E, Howarth RM, Boczonadi V, et al. Salbutamol modifies the neuromuscular junction in a mouse model of ColQ myasthenic syndrome. Hum Mol Genet. 2019;28(14):2339–51. doi:10.1093/hmg/ddz059

23. O’Connor E, Töpf A, Müller JS, Cox D, Evangelista T, Colomer J, et al. Identification of mutations in the MYO9A gene in patients with congenital myasthenic syndrome. Brain. 2016;139(8):2143–53. doi:10.1093/brain/aww130 PubMed PMID: 27259756.

24. McMacken G, Abicht A, Evangelista T, Spendiff S, Lochmüller H. The Increasing Genetic and Phenotypical Diversity of Congenital Myasthenic Syndromes. Neuropediatrics. 2017;48(4):294–308. doi:10.1055/s-0037-1602832 PubMed PMID: 28505670.

25. Natera-de Benito D, Töpf A, Vilchez JJ, González-Quereda L, Domínguez-Carral J, Díaz-Manera J, et al. Molecular characterization of congenital myasthenic syndromes in Spain. Neuromuscular Disorders. 2017 Dec 1;27(12):1087–98. doi:10.1016/j.nmd.2017.08.003

26. O’Connor E, Phan V, Cordts I, Cairns G, Hettwer S, Cox D, et al. MYO9A deficiency in motor neurons is associated with reduced neuromuscular agrin secretion. Vol. 27. 2018;27(8):1434–46. doi:10.1093/hmg/ddy054

27. Wen H, Brehm P. Paired patch clamp recordings from motor-neuron and target skeletal muscle in zebrafish. Journal of Visualized Experiments. 2010;(45). doi:10.3791/2351

28. Wen H, Eckenstein K, Weihrauch V, Stigloher C, Brehm P. Primary and secondary motoneurons use different calcium channel types to control escape and swimming behaviors in zebrafish. Proc Natl Acad Sci U S A. 2020 Oct 20;117(42):26429–37. doi:10.1073/PNAS.2015866117;WGROUP:STRING:PUBLICATION PubMed PMID: 33020266.

29. Wen H, Linhoff MW, Hubbard JM, Nelson NR, Stensland D, Dallman J, et al. Zebrafish Calls for Reinterpretation for the Roles of P/Q Calcium Channels in Neuromuscular Transmission. Journal of Neuroscience. 2013 Apr 24;33(17):7384–92. doi:10.1523/JNEUROSCI.5839-12.2013 PubMed PMID: 23616544.

30. Wen H, Hubbard JM, Rakela B, Linhoff MW, Mandel G, Brehm P. Synchronous and asynchronous modes of synaptic transmission utilize different calcium sources. Elife. 2013 Dec 24;2013(2). doi:10.7554/ELIFE.01206 PubMed PMID: 24368731.

31. Wen H, McGinley MJ, Mandel G, Brehm P. Nonequivalent release sites govern synaptic depression. Proceedings of the National Academy of Sciences. 2016 Jan 19;113(3):E378–86. doi:10.1073/PNAS.1523671113 PubMed PMID: 26715759.

32. Wyart C, Bene F Del, Warp E, Scott EK, Trauner D, Baier H, et al. Optogenetic dissection of a behavioural module in the vertebrate spinal cord. Nature. 2009 Sep 17;461(7262):407–10. doi:10.1038/nature08323 PubMed PMID: 19759620.

33. Antinucci P, Dumitrescu AS, Deleuze C, Morley HJ, Leung K, Hagley T, et al. A calibrated optogenetic toolbox of stable zebrafish opsin lines. Elife. 2020 Mar 1;9. doi:10.7554/ELIFE.54937 PubMed PMID: 32216873.

34. Whittaker RG, Herrmann DN, Bansagi B, Hasan BAS, Lofra RM, Logigian EL, et al. Electrophysiologic features of SYT2 mutations causing a treatable neuromuscular syndrome. Neurology. 2015 Dec 1;85(22):1964–71. doi:10.1212/WNL.0000000000002185 PubMed PMID: 26519543.

35. Drapeau P, Ali DW, Buss RR, Saint-Amant L. In vivo recording from identifiable neurons of the locomotor network in the developing zebrafish. J Neurosci Methods. 1999 Apr 1;88(1):1–13. doi:10.1016/S0165-0270(99)00008-4 PubMed PMID: 10379574.

36. Ham AS, Lin S, Tse A, Thürkauf M, McGowan TJ, Jörin L, et al. Single-nuclei sequencing of skeletal muscle reveals subsynaptic-specific transcripts involved in neuromuscular junction maintenance. Nature Communications . 2025 Dec 1;16(1):1–19. doi:10.1038/S41467-025-57487-1;TECHMETA PubMed PMID: 40044687.

37. Hui T, Jing H, Lai X. Neuromuscular junction-specific genes screening by deep RNA-seq analysis. Cell Biosci. 2021 Dec 1;11(1):1–15. doi:10.1186/S13578-021-00590-9/FIGURES/8

38. Bedell VM, Westcot SE, Ekker SC. Lessons from morpholino-based screening in zebrafish. Brief Funct Genomics. 2011 Jul;10(4):181–8. doi:10.1093/bfgp/elr021 PubMed PMID: 21746693.

39. Nicklin J, Karni Y, Wiles CM. Shoulder abduction fatiguability. J Neurol Neurosurg Psychiatry. 1987;50(4):423–7. doi:10.1136/jnnp.50.4.423 PubMed PMID: 3585353.

40. Kaeser PS, Regehr WG. Molecular Mechanisms for Synchronous, Asynchronous, and Spontaneous Neurotransmitter Release. Annu Rev Physiol. 2013 Feb;76:333. doi:10.1146/ANNUREV-PHYSIOL-021113-170338 PubMed PMID: 24274737.

41. Szklarczyk D, Kirsch R, Koutrouli M, Nastou K, Mehryary F, Hachilif R, et al. The STRING database in 2023: protein–protein association networks and functional enrichment analyses for any sequenced genome of interest. Nucleic Acids Res. 2023 Jan 6;51(D1):D638–46. doi:10.1093/nar/gkac1000 PubMed PMID: 36370105.

42. Krämer A, Green J, Pollard J, Tugendreich S. Causal analysis approaches in Ingenuity Pathway Analysis. Bioinformatics. 2014 Feb 15;30(4):523–30. doi:10.1093/bioinformatics/btt703 PubMed PMID: 24336805.

43. Engel AG, Selcen D, Shen XM, Milone M, Harper CM. Loss of MUNC13-1 function causes microcephaly, cortical hyperexcitability, and fatal myasthenia. Neurol Genet. 2016;2(5):e105. doi:10.1212/NXG.0000000000000105;WGROUP:STRING:PUBLICATION

44. Shen XM, Selcen D, Brengman J, Engel AG. Mutant SNAP25B causes myasthenia, cortical hyperexcitability, ataxia, and intellectual disability. Neurology. 2014 Dec 9;83(24):2247–55. doi:10.1212/WNL.0000000000001079 PubMed PMID: 25381298.

45. Pugliese A, Holland SH, Rodolico C, Lochmüller H, Spendiff S. Presynaptic Congenital Myasthenic Syndromes: Understanding Clinical Phenotypes through In vivo Models. J Neuromuscul Dis. 2023 May 19;10(5):731–59. doi:10.3233/JND-221646 PubMed PMID: 37212067.

46. Genç Ö, Kochubey O, Toonen RF, Verhage M, Schneggenburger R. Munc18-1 is a dynamically regulated PKC target during short-term enhancement of transmitter release. Elife. 2014 Feb 11;2014(3):e01715. doi:10.7554/ELIFE.01715.001 PubMed PMID: 24520164.

47. Rizo J, Südhof TC. Snares and munc18 in synaptic vesicle fusion. Nature Reviews Neuroscience. European Association for Cardio-Thoracic Surgery; 2002. p. 641–53. doi:10.1038/nrn898

48. Südhof TC. Neurotransmitter Release: The Last Millisecond in the Life of a Synaptic Vesicle. Neuron. 2013 Oct 30;80(3):10.1016/j.neuron.2013.10.022. doi:10.1016/J.NEURON.2013.10.022 PubMed PMID: 24183019.

49. Pennuto M, Bonanomi D, Benfenati F, Valtorta F. Synaptophysin I Controls the Targeting of VAMP2/Synaptobrevin II to Synaptic Vesicles. 2003 Oct 3;14(12):4909–19. doi:10.1091/mbc.E03-06-0380 PubMed PMID: 14528015.

50. Pang ZP, Melicoff E, Padgett D, Liu Y, Teich AF, Dickey BF, et al. Synaptotagmin-2 Is Essential for Survival and Contributes to Ca2+ Triggering of Neurotransmitter Release in Central and Neuromuscular Synapses. Journal of Neuroscience. 2006 Dec 27;26(52):13493–504. doi:10.1523/JNEUROSCI.3519-06.2006 PubMed PMID: 17192432.

51. Melicoff E, Sansores-Garcia L, Gomez A, Moreira DC, Datta P, Thakur P, et al. Synaptotagmin-2 Controls Regulated Exocytosis but Not Other Secretory Responses of Mast Cells. J Biol Chem. 2009 Jul 17;284(29):19445. doi:10.1074/JBC.M109.002550 PubMed PMID: 19473977.

52. Bouhours B, Gjoni E, Kochubey O, Schneggenburger R. Synaptotagmin2 (Syt2) Drives Fast Release Redundantly with Syt1 at the Output Synapses of Parvalbumin-Expressing Inhibitory Neurons. Journal of Neuroscience. 2017 Apr 26;37(17):4604–17. doi:10.1523/JNEUROSCI.3736-16.2017 PubMed PMID: 28363983.

53. Thaker H, Zhang J, Miyashita SI, Cristofaro V, Park SH, Hashemi-Gheinani A, et al. Knockin mouse models demonstrate differential contributions of synaptotagmin-1 and -2 as receptors for botulinum neurotoxins. PLoS Pathog. 2021 Oct 1;17(10):e1009994. doi:10.1371/JOURNAL.PPAT.1009994 PubMed PMID: 34662366.

54. Chen C, Arai I, Satterfield R, Young SM, Jonas P. Synaptotagmin 2 Is the Fast Ca2+ Sensor at a Central Inhibitory Synapse. Cell Rep. 2017 Jan 17;18(3):723–36. doi:10.1016/J.CELREP.2016.12.067 PubMed PMID: 28099850.

55. Tuvim MJ, Mospan AR, Burns KA, Chua M, Mohler PJ, Melicoff E, et al. Synaptotagmin 2 Couples Mucin Granule Exocytosis to Ca2+ Signaling from Endoplasmic Reticulum. J Biol Chem. 2009 Apr 10;284(15):9781. doi:10.1074/JBC.M807849200 PubMed PMID: 19208631.

56. Pang ZP, Sun J, Rizo J, Maximov A, Südhof TC. Genetic analysis of synaptotagmin 2 in spontaneous and Ca2+-triggered neurotransmitter release. EMBO J. 2006 May 17;25(10):2039. doi:10.1038/SJ.EMBOJ.7601103 PubMed PMID: 16642042.

57. Westerfield M, McMurray J V., Eisen JS. Identified motoneurons and their innervation of axial muscles in the zebrafish. Journal of Neuroscience. 1986;6(8):2267–77. doi:10.1523/jneurosci.06-08-02267.1986 PubMed PMID: 3746409.

58. Bello-Rojas S, Istrate AE, Kishore S, McLean DL. Central and peripheral innervation patterns of defined axial motor units in larval zebrafish. Journal of Comparative Neurology. 2019 Oct 15;527(15):2557–72. doi:10.1002/CNE.24689;ISSUE:ISSUE:DOI PubMed PMID: 30919953.

59. Srinivasan SS, Maimon BE, Diaz M, Song H, Herr HM. Closed-loop functional optogenetic stimulation. Nature Communications 2018 9:1. 2018 Dec 13;9(1):5303-. doi:10.1038/s41467-018-07721-w PubMed PMID: 30546051.

60. Zamani A, Sakuragi S, Ishizuka T, Yawo H. Kinetic characteristics of chimeric channelrhodopsins imsplicate the molecular identity involved in desensitization. Biophys Physicobiol. 2017;14:13. doi:10.2142/BIOPHYSICO.14.0_13 PubMed PMID: 28409086.

61. Pang ZP, Xu W, Cao P, Südhof TC. Calmodulin Suppresses Synaptotagmin-2 Transcription in Cortical Neurons,. Journal of Biological Chemistry. 2010 Oct 29;285(44):33930–9. doi:10.1074/JBC.M110.150151 PubMed PMID: 20729199.

62. Liao W, Elfrink K, Bähler M. Head of Myosin IX Binds Calmodulin and Moves Processively toward the Plus-end of Actin Filaments. Journal of Biological Chemistry. 2010 Aug 6;285(32):24933–42. doi:10.1074/jbc.M110.101105 PubMed PMID: 20538589.

63. Junge HJ, Rhee JS, Jahn O, Varoqueaux F, Spiess J, Waxham MN, et al. Calmodulin and Munc13 Form a Ca2+ Sensor/Effector Complex that Controls Short-Term Synaptic Plasticity. Cell. 2004 Aug 6;118(3):389–401. doi:10.1016/j.cell.2004.06.029 PubMed PMID: 15294163.

64. Tasnim M, Wahlquist P, Hill JT. Zebrafish: unraveling genetic complexity through duplicated genes. Dev Genes Evol. 2024 Dec 1;234(2):99. doi:10.1007/S00427-024-00720-6 PubMed PMID: 39079985.

